# sciCSR infers B cell state transition and predicts class-switch recombination dynamics using single-cell transcriptomic data

**DOI:** 10.1101/2023.02.02.526789

**Authors:** Joseph CF Ng, Guillem Montamat Garcia, Alexander T Stewart, Paul Blair, Deborah K Dunn-Walters, Claudia Mauri, Franca Fraternali

## Abstract

Class-switch recombination (CSR) is an integral part of B cell maturation. Steady-state analyses of isotype distribution (e.g. B cell receptor [BCR] repertoire analysis of snapshots during an immune response) do not directly measure CSR dynamics, which is crucial in understanding how B cell maturation is regulated across time. We present sciCSR (pronounced ‘scissor’, single-cell inference of class switch recombination), a computational pipeline which analyses CSR events and dynamics of B cells from single-cell RNA-sequencing (scRNA-seq) experiments. sciCSR re-analyses transcriptomic sequence alignments to differentiate productive heavy-chain immunoglobulin transcripts from germline “sterile” transcripts. From a snapshot of B cell scRNA-seq data, a Markov state model is built by the pipeline to infer the dynamics and direction of CSR. Applying sciCSR on SARS-CoV-2 vaccination time-course scRNA-seq data, we observe that sciCSR predicts, using data from an earlier timepoint in the collected time-course, the isotype distribution of BCR repertoires of subsequent timepoints with high accuracy (cosine similarity ∼ 0.9). sciCSR also recapitulates CSR patterns in mouse models where B cell maturation was perturbed using gene knockouts. sciCSR infers cell state transitions using processes specific to B cells, identifies transitions which are often missed by conventional RNA velocity analyses, and can reveal insights into the regulation of CSR and the dynamics of B cell maturation during an immune response.

## Introduction

B cells are the main drivers of the humoral response in developing protection against infectious diseases. An understanding of how this process is regulated over time is crucial to evaluate the quality of the antibodies produced and, in turn, the effectiveness of vaccination and therapeutic strategies^1-3^. B cells mature from a naïve state to acquire memory against the antigen and differentiate into antibody-producing plasma cells^1, 3, 4^. Traditionally, the dynamics of B cell maturation is investigated by isolating different B cell subsets based on their surface protein expression, and comparing the proportion of these subsets before and after antigen exposure, or through a more detailed time-course^59^. Such subsets are typically distinct cell states, and one would require additional experimental evidence to probe the molecular details of how cells transition from one state into another. Single-cell profiling offers rich descriptions of the transcriptomic features of B cell states and how they are altered during B cell maturation^10–12^. In combination with B cell receptor (BCR) repertoires, these data have yielded great insights into the heterogeneity and functional relevance of different B cell subsets in health and disease^10,11, 13–16^.

In the past few years a plethora of computational methods have been created to infer the dynamics of cell state transitions from single-cell RNA sequencing (scRNA-seq) data. Typically they fall into two categories: the first belongs to a large family of methods which infer “pseudotime” ordering of cells^17, 18^, sometimes requiring user input of the likely start and end states; the second corresponds to exploiting the balance between unspliced and spliced reads to infer what is called “RNA velocity”^19, 20^. “Pseudotime” ordering is derived from “trajectories” of cell differentiation fitted to the data by imposing directionality, often using prior knowledge, onto a transcriptional similarity network of cells. As such, cells can be ordered along a “pseudotime” axis often understood to encode the differentiation potential of cells^21, 22^. Pseudotime can be informed by the underlying data even in the absence of an actual time-course in sample collection. The second category of tools which describe cell transition dynamics estimate “RNA velocity” by considering the splicing kinetics for genes^19, 20^. The observation of nascent, unspliced mRNA and its mature, spliced counterparts across single cells allows one to extrapolate the future state of the system given its current state^19^. Recent years have seen methodological development in the estimation of RNA velocity, in accounting for variations of splicing rates and expression levels across genes to improve the reconstruction of cell differentiation trajectories^20, 23–25^. This accompanies experimental approaches to capture the time component of cellular development, e.g. lineage tracing^26–28^ and metabolite labelling^29, 30^ approaches coupled to scRNA-seq data generation. Computationally, methods such as CellRank^31^ build on both RNA velocity and pseudotime methods, to fit statistical models which describe the overall dynamics of cell state transitions observed in the data. These methods, although successful in experimental validation, are designed and tested on use cases in developmental biology, where typically both the progenitor(s) and mature cell state(s) are transcriptionally distinct and well-defined. Haematopoiesis has been studied with scRNA-seq, including the application of RNA velocity and pseudotime ordering tools, to some degree of success^17, 29^. On the other hand, it is posited that samples such as peripheral blood mononuclear cells (PBMCs) are cell types with already equilibrated RNA metabolism, lacking the dynamic information required for RNA velocity analysis. This limitation results in noisy velocity profiles and erroneous inference of transitions^32^. Whilst computational approaches begin to offer diagnostic approaches to determine the suitability of RNA velocity analysis on these datasets^24^, a viable alternative to study transitions in scRNA-seq datasets of immune cell types is lacking.

This raises methodological questions on how to improve the approaches to study transition dynamics in immunological systems, especially mature cell types such as B cells in circulation and in secondary lymphoid tissues. Notably, B cells continue to mature after exiting the bone marrow by somatic hypermutation (SHM) and class-switch recombination (CSR)^33-36^ to optimise the B cell receptor (BCR) for function. SHM targets the variable (V), diversity (D) and joining (J) gene segments and introduces mutations catalysed by activation-induced cytidine deaminase (AID)^34,36–38^ (**Figure 1**a, left) to optimise antigen binding. On the other hand, CSR alters the constant (C) region to adapt the BCR to function in different immune challenges and tissues contexts. CSR also depends on AID, which recognises “switch” genomic regions 5’ to each C gene enriched in motives such as 5’-AGCT-3’^35, 39-42^ and catalyses deamination^35, 40, 43^ (**Figure 1**a, right). The repair of these mutational events entails DNA recombination that brings the immunoglobulin variable region proximal to a downstream C gene which, when transcribed and translated, would encode an immunoglobulin molecule switched to this isotype^44, 45^. AID is targeted to the desired downstream C gene in part by transcriptional activity originated from specific combination of molecular signals; such transcription initiates at positions immediately 5’ to the target C gene and, as such, lacks the VDJ gene segments to encode a fully functional immunoglobulin heavy chain^35, 46–49^. The production of these germline or “sterile” transcripts therefore signifies that CSR events are predisposed to occur^35,50-52^ (**Figure 1**a, right). Theoretically, both CSR and SHM would leave transcriptomic footprints (BCR sequence substitutions for SHM; sterile transcripts for CSR) which can be captured in scRNA-seq and scBCR-seq (i.e. profiling of the BCR repertoire barcoded with originating cell and molecule identifiers) data (**Figure 1**b,c). Neither RNA velocity or pseudotime methods have access to CSR and SHM, but it is these processes which are of interest in describing B cell maturation and the function of the resultant antibodies secreted during an immune response^2, 3, 35^. Extracting signals of CSR and SHM from the data, and utilising them to infer B cell transitions, would complement existing velocity/pseudotime methods and yield a more faithful reconstruction of B cell maturation dynamics from single-cell transcriptomic data.

**Figure 1.**
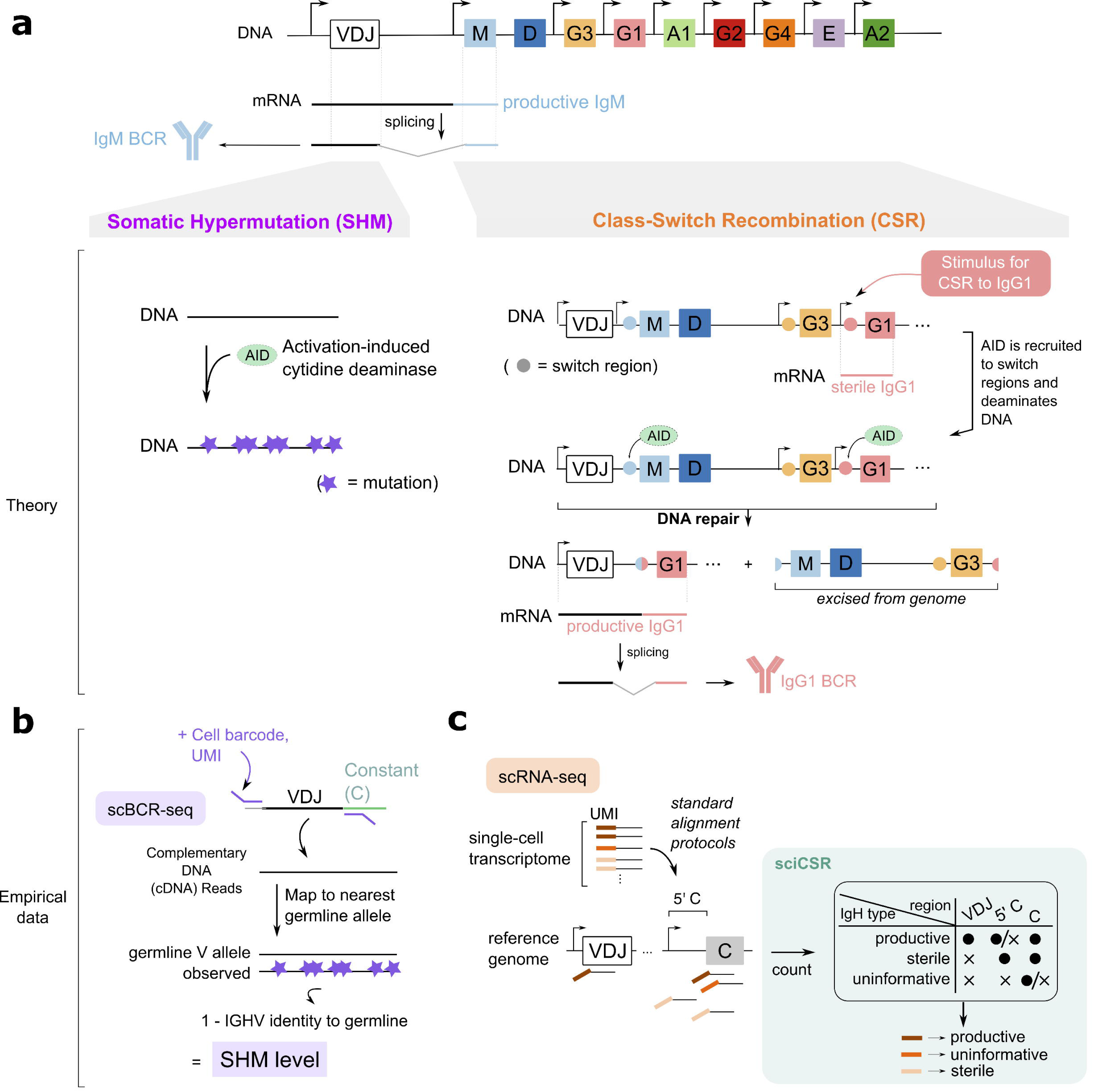
Overview of the theories of somatic hypermutation (SHM) and class-switch recombination (CSR) and their footprints in empirical data. (a) *(top)* Schematic of the immunoglobulin heavy-chain genomic locus. In Naïve B cells, transcription of the VDJ segment through to the IgM gene generates a productive IgM transcript which encodes the heavy chain of the IgM BCR protein. *(middle)* SHM acts on t**he** VDJ segment, where mutations are introduced by activation-induced cytidine deaminase (AID). CSR is a DNA recombination process involving the constant (C) region genes: each C gene (except IgD) is a discrete transcriptional unit controlled by its own promoter. Stimuli for switching towards a specific isotype (here illustrated as IgG1) would activate the transcription of a “sterile” transcript which do not contain the VDJ segment. It is thought that AID is recruited to positions 5’ to the C gene coding segments (“switch region”), to initiate a series of DNA damage and repair events. As a result, the recombination of the upstream VDJ region with the segment encoding the downstream switched isotype would subsequently produce a “productive” transcript encoding a full-length heavy chain. The DNA fragment encoding the previous isotype(s) is excised from the genome. (b-c) Both SHM and CSR can be observed in sequencing datasets: SHM can be analysed in scBCR-seq (b), by comparing the observed VDJ sequences with the nearest germline allele and enumerating mutations. CSR can be analysed in scRNA-seq (c); sciCSR provides functionalities to reconstitute each heavy-chain transcript as productive or sterile, based on whether reads mapping to the VDJ, C, or 5’ C (i.e. region 5’ to the C region coding segment) regions can be observed for each transcript.

Here we introduce sciCSR (pronounced “scissor”, acronym for “single-cell inference of class switch recombination”), a computational tool designed to infer state transitions between mature B cell states and predict the direction of CSR in B cells assayed using scRNA-seq. Motivated by the need to faithfully capture B cell state transitions, sciCSR implements routines which extract B-cell-native information, such as CSR and SHM, from scRNA-seq data and, if available, BCR repertoire of the same population of B cells. This information is used as input to CellRank to infer transition probabilities between cells. We further applied transition path theory (TPT) onto this transition matrix to analyse the dynamics of transitions between cell clusters; this method, popularised in the analysis of conformational state transitions in molecular dynamics simulation data^53–55^, describes transitions by considering an ensemble of possible paths governed by the given transition matrix and, as such, allows for more flexible analysis of transitions beyond visualisation of arrows depicting velocity streams which are typical in RNA velocity analysis. We validated sciCSR using immunisation and gene knockout studies, and showed that sciCSR could recover BCR isotype distributions at the steady-state and capture CSR dynamics which could be subsequently verified over time-courses.

## Results

### Extracting class-switch recombination signals from single-cell RNA-sequencing data of B cells

We hypothesise that the use of information native to B cells, namely class-switch recombination (CSR) and somatic hypermutation (SHM), would improve the inference of cell state transitions in mature B cells as observed in scRNA-seq data. SHM level can be easily retrieved from BCR sequencing data of single cells which is routinely obtained in parallel to transcriptome-wide single-cell profiling: SHM is inversely related to the sequence identity between the observed BCR sequence and the corresponding germline immunoglobulin gene^57-60^ (**Figure 1**b). The detection of CSR, on the other hand, is less trivial at the single-cell level: CSR is more easily characterized by either comparing the expression level of different immunoglobulin heavy chain (IgH) isotypes at the transcript or protein level for a cell population^61–63^, or by studying BCR clonotypes which comprise sequences of different isotypes^57, 64, 65^. In sciCSR a series of routines has been implemented to distinguish productive and sterile IgH transcripts from scRNA-seq data, by enumerating mRNA molecules (identifiable by observing combinations of cell barcode and unique molecular identifier (UMI)) with reads mapping to the VDJ, 5’ C or C regions in the immunoglobulin heavy-chain (IgH) genomic locus (**Figure 1**c). Molecules with insufficient evidence to be classified as productive or sterile (e.g. if only one read mapped to the C exonic regions is found) are labelled as ‘uninformative’. This quantification complements conventional data processing workflow: in standard genome annotations, individual immunoglobulin V, D, J and C genes are treated as separate gene entities; reads which are 5’ of C genes would therefore be considered as intergenic reads typically discarded in transcriptomic data analysis workflows.

We first asked whether these 5’ C reads could be detected in B cell scRNA-seq data. We previously sorted mature, circulating human B cells into different phenotypically defined subsets and generated scRNA-seq libraries^10^ using the 5’ 10X Genomics protocol; **Figure 2**a shows the distribution of sequencing reads across the IgH genomic locus. We observed reads mapped to regions 5’ to *IGHG1* and *IGHG2* in unswitched B cell subsets (CD19+IgD+CD27’CD10’ naïve B cells and CD19+IgD+CD27+ IgM memory cells), and notably with peaks at the transcription start sites (TSS). While these reads could be consistent with either genuine sterile IgH transcripts, or productive IgH transcripts where the intron between the variable and constant regions is yet to be spliced, we reason that the latter case is unlikely given the scRNA-seq library preparation protocol enriches mRNA at the 5’ end: in our data we observe that reads that were concentrated at typically around 600 base-pairs downstream of TSS (when inspecting single-exon transcripts e.g. *JUNB, RHOB,* to eliminate splicing effects on the read distributions, see Supplementary Figure S2), and the IgH read peaks in **Figure 2**a would have been around 1-2 kilobase downstream of the TSS of a productive IgH transcript, thus discounting the likelihood that IgH transcripts bearing the VDJ segments were the sources of these 5’ C reads.

**Figure 2.**
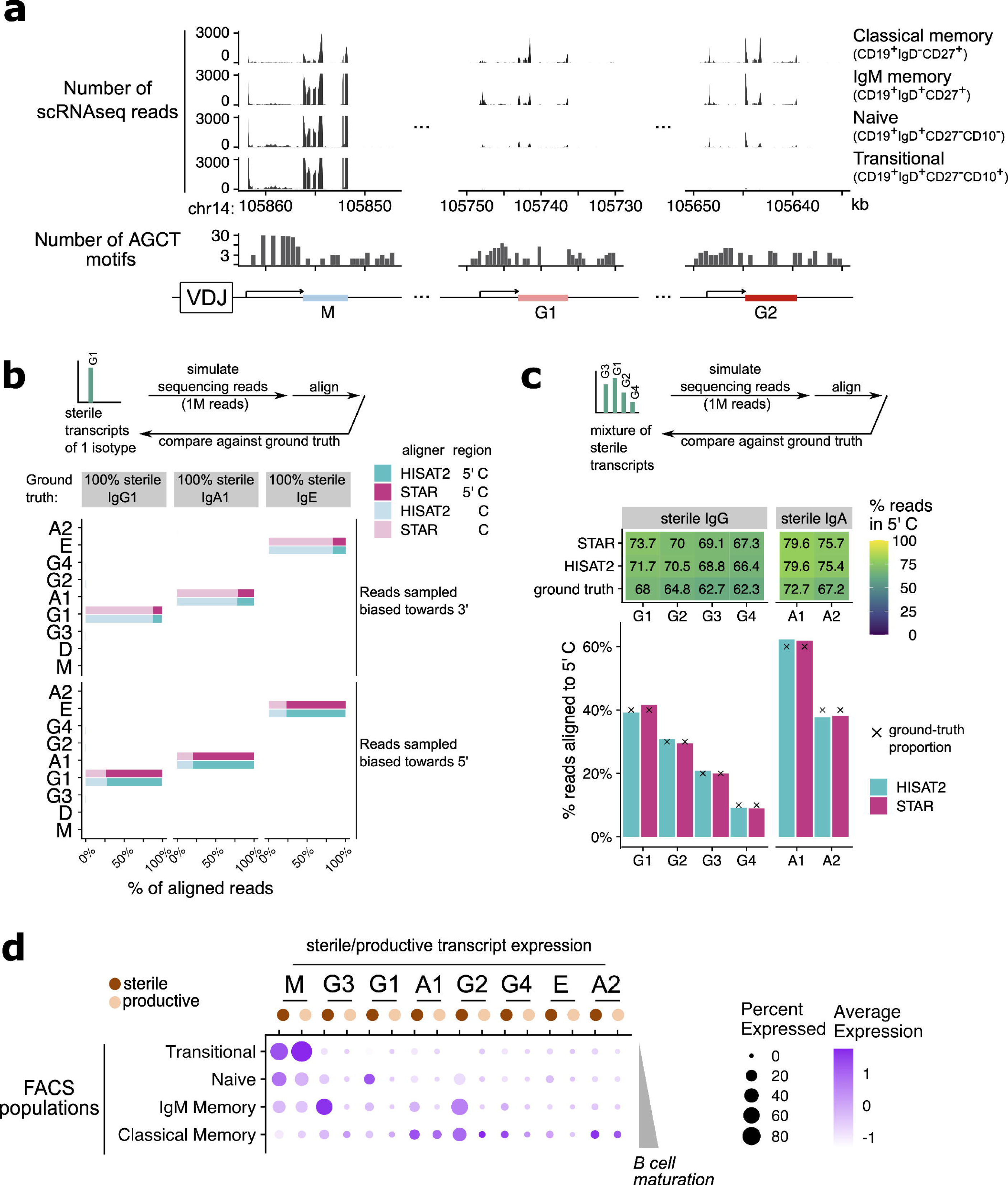
sciCSR reconstitutes productive and sterile IgH transcription levels in different B cell subsets. (a) Histograms of scRNA-seq reads for mature human B cells from peripheral blood FACS-sorted into different B cell subsets (data from Stewart et al). Only reads mapped to IgM, IgG1 and IgG2 are shown here; see Supplementary Figure S1 for the complete IgH locus. Notice the IgH coding sequences are on the minus strand of human chromosome 14; the horizontal axis is depicted in reverse. (b) Alignment of simulated reads (using the polyester package), sampled from IgG1, IgA1 and IgE which are mapped to the 5’ C (light shade) or the C exons (dark shade), using either HISAT2 (teal) or STAR (magenta). Reads sampling are biased either to the 3’ (top panel) or the 5’ (bottom) end of transcripts to mimic typical scRNA-seq library preparation protocols. (c) Alignment of polyester-simulated reads sampled from sterile transcripts of a mixture of IgG (left) or IgA (right) subtypes. The heatmap at the top depicts the proportion of sampled reads mapped to the 5’ C region which is informative of indicating sterile IgH. The bar plots depict the proportions of reads aligned to each isotype using HISAT2 (teal) or STAR (magenta). The ground-truth proportions were noted by crosses (X). (d) Dotplot depicting productive and sterile transcription level recovered using sciCSR for the Stewart et al dataset shown in (a). Dot size corresponds to the proportion of cells with positive expression while colour intensity represents expression level.

Applying the sciCSR routine depicted in **Figure 1** to simulated reads sampled from the human IgH genomic locus (see Materials and Methods), we confirm that commonly used RNA read aligners STAR^66^ (which is used in the 10X cellranger pre-processing workflow) and HISAT2^67^ can accurately identify the isotype corresponding to sterile IgH transcripts (**Figure 2**b); notably, these aligners are precise in identifying the exact IgG and IgA subtypes, supporting the direct use of aligned reads generated by these data pre-processing pipelines to analyse productive/sterile IgH transcription in sciCSR. In further support of this, we find that both aligners can accurately recover the composition of mixtures of sterile IgG and IgA transcripts simulated with fixed proportion of each subtype (**Figure 2**c). The major requirement for sciCSR is the adoption of a 5’ enrichment protocol in the library preparation step, as reads biased to the 3’ end do not capture the 5’ C region necessary to define sterile transcripts (**Figure 2**b). The simulation results suggest that given this prerequisite is fulfilled, the aligned reads generated by commonly used aligners in scRNA-seq data pre-processing workflows can be directly used to analyse sterile IgH levels using sciCSR. The sciCSR workflow re-analyses the pool of immunoglobulin transcripts from a single mature B cell, and extracts information about both its current state (the IgH isotype it currently produces) and its immediate future^50^, ideal as a basis to build models which describe B cell transition dynamics.

### The sciCSR pipeline

The routine above generates count matrices just like the standard transcriptome-wide count data typically used for downstream scRNA-seq analysis, such as differential expression and visualisation of expression levels. For example, **Figure 2**d shows the sterile and productive IgM transcription levels in the Stewart *et al* dataset^10^ visualised in **Figure 2**a. This captures, along the B cell maturation process, the switches away from IgM, the rise of sterile transcription (most notably IgG subtypes) in the IgM memory cells and ultimately productive transcription of IgG and IgA subtypes in the classical memory subset. We do, however, note that these counts, especially those corresponding to productive IgH, are fairly sparse (Supplementary Figure S3) given their definition requires at least 2 reads mapped for each molecule.

To derive metrics for inferring cellular transition dynamics, it is advisable to leverage signals from productive and sterile transcripts of all IgH C genes collectively for robustness; ideally, metrics which summarise CSR status should also be comparable across datasets, such that they are robust to different data integration protocols commonly used to aggregate scRNA-seq datasets. In sciCSR we address this by defining “isotype signatures” using non-negative matrix factorisation^68^ (NMF) over the productive and sterile transcripts of all isotypes from reference data (**Figure 3**a), and using these signatures to score the CSR status of cells. These “isotype signatures” describe the expression level of all IgH productive and sterile transcripts for naïve/memory B cell states. The other output of the NMF analysis is a weighting specific for each B cell in the data, which are scores indicating its resemblance to naïve/memory state based on the observed productive/sterile IgH counts (**Figure 3**a). The weighting for the naïve state for each cell is taken to define a metric which we term “CSR potential”; this metric orders cells from naïve to isotype-switched state (which would typically comprise both classical memory B cells and switched plasmablast/plasma cells). Visualising the distribution of “CSR potential” in real data, we observe that this metric orders B cells correctly by their maturation stages (**Figure 3**b), which can be cross-referenced with changes in productive and sterile transcription levels (**Figure 3**c). To generate CSR potentials which can be compared across different conditions, we apply NMF on reference atlases of mouse and human B cells^10, 11, 13, 14^ covering a range of cell states present in germinal centre (GC) and circulating B cells, to learn isotype signatures separately for each species. These pre-trained signatures can be used directly to obtain the CSR potential from new, user-supplied datasets (**Figure 3**d). If parallel libraries of BCR sequences (hereafter “scBCR-seq”) are available, sciCSR can also calculate the SHM level of each cell, by computing the quantity (1 – percentage identity to germline IgH V gene). Both CSR potential and SHM frequency can be interpreted effectively as pseudotime ordering of B cells defined based on these processes which are biologically relevant to B cell maturation. These metrics are used as input to the CellRank algorithm^31^ to impose directionality onto the cell-cell *k*-nearest neighbour (kNN) network, to derive a transition matrix which describes the probability of transition between cells (**Figure 3**d). We can directly compare the inference results using CSR/SHM information to conventional CellRank analyses which uses RNA velocity to bias the kNN network.

**Figure 3.**
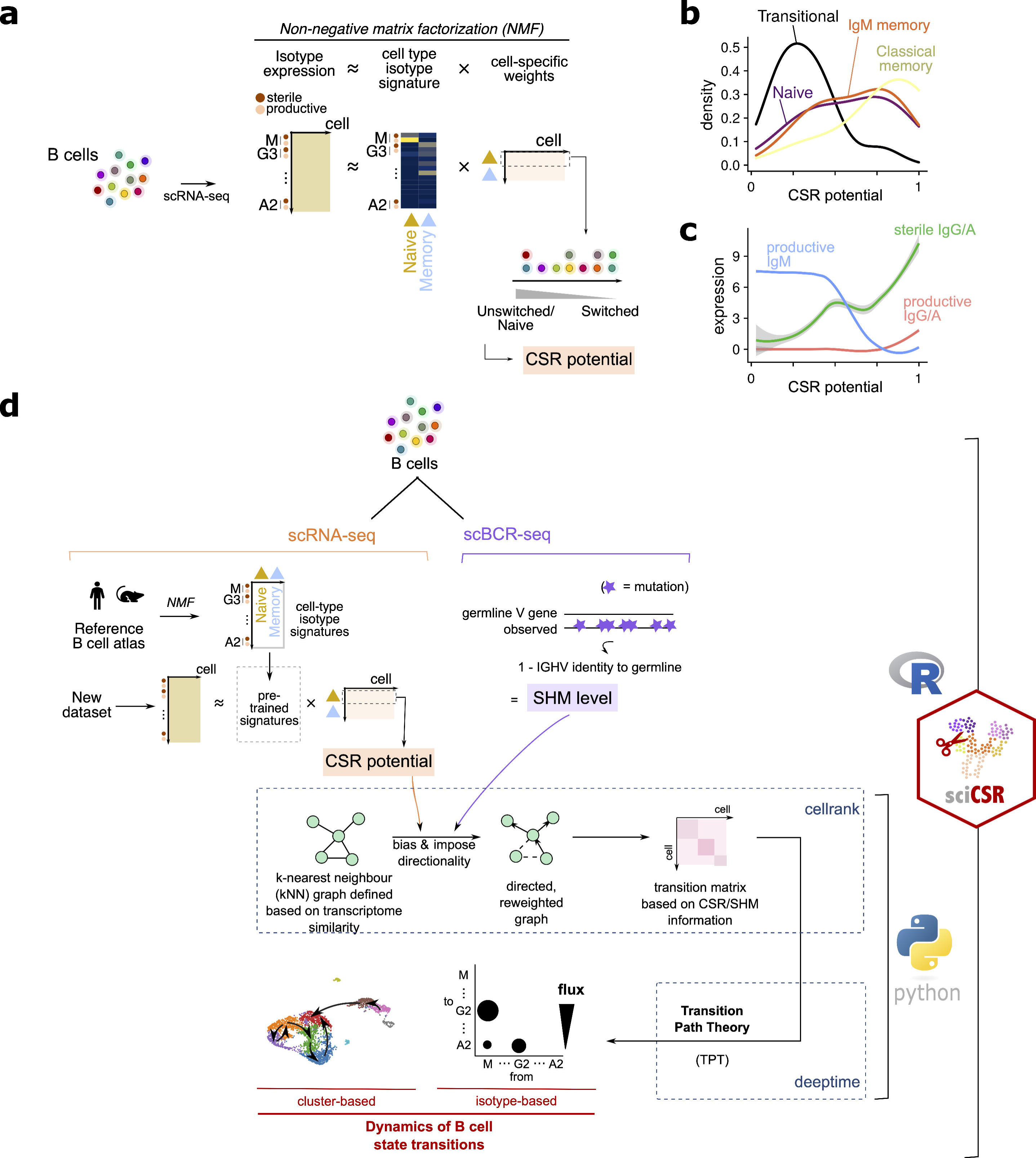
The sciCSR pipeline to infer B cell state transitions. (a) Reference B cell atlases from mouse and human were subjected to non-negative matrix factorization (NMF) to extract isotype signatures (see main text) which describe naïve/memory B cell subsets. The naïve signature was used to rank the cells by their maturation status, to generate a score termed ‘CSR potential’. (b) Distribution of CSR potential score for B cell subsets in the Stewart et al. dataset shown in Figures 2a,d. (c) Expression level of different productive and sterile IgH transcripts along the axis of the inferred CSR potentials in the Stewart et al. dataset. (d) Schematic to illustrate the sciCSR workflow for inferring state transitions. sciCSR analyses scRNA-seq and (if available) scBCR-seq data to rank cells by their CSR/SHM levels, and usea this ranking as inputs to CellRank to construct Markov state models, subsequently analysed using Transition Path Theory (TPT), which describes the transition dynamics of cell-types/isotypes. Pre-trained NMF signatures from the reference atlases (see panel a) were used to deconvolute user-supplied dataset. If scBCR-seq data is available, SHM level is calculated by considering mutations on IgH V gene sequences. These CSR/SHM-based rankings are used to bias the cell *ƙ*-nearest neighbour (kNN) network to impose directionality to describe cell state transitions. See main text and Materials and Methods for further details.

CellRank automatically aggregates individual cells into “macrostates” which share similar transitional behaviours and detects the start and end points of the transition pathway by examining the properties of the cell-cell transition matrix. In our application to B cells, we require a more flexible analytical framework to accommodate use cases where (i) cell states are defined not solely by transcriptome-wide features, but rather by using other criteria of interest (e.g. in CSR, where “states” are defined based on the productive IgH isotype of individual B cells, but not necessarily its overall transcriptomic profile), and; (ii) multiple possible traversal pathways are possible and expected (e.g. in CSR, one could switch isotype in a stepwise manner along the IgH genomic locus, or jump directly between isotypes whilst skipping those located physically between the source and destination isotype C genes). To address these challenges, sciCSR incorporates the following innovations in analysing cell transition dynamics based on scRNA-seq data: first, we introduce Transition Path Theory (TPT) to model dynamics implied in the transitions between the cell groupings (**Figure 3**d). TPT is used in physics and chemistry to understand how the potential energy landscape is explored in chemical reactions or in conformational changes^56, 69, 70^, and popularised in biology to analyse biomolecular dynamics simulation data^53^”^55^. Instead of nominating a single pathway to navigate the state landscape, TPT describes the statistical properties of the ensemble of possible transition paths given the transition matrix; state transitions are characterised by the amount of flux between states, and a list of possible traversal pathways are nominated and ranked by their likelihoods given the flux estimates. The inclusion of multiple traversals provides flexibility in modelling transitions more relevant to contexts such as stepwise versus direct CSR events. Second, to robustly identify unidirectional transitions such as CSR, we reason that insignificant fluxes should have magnitudes comparable to “null” transition models defined by reshuffling columns of the transition matrix; sciCSR carries out random shuffling of the transition matrix and fits null TPTs, from which p-values are calculated to evaluate the relevance of fluxes inferred from the data, in order to assist the interpretation of TPT fluxes and down-weigh random transitions. Finally, sciCSR groups cells either by user-defined cluster labels or by the BCR isotypes they express. This allows for the inference of transitions at both the cell cluster level to analyse the trajectories of B cell subsets in the data, or infer CSR which are directly testable by comparing the inferred fluxes against the isotype distributions observed in BCR repertoire sequencing data (**Figure 3**d).

sciCSR is implemented as a R package which manipulates Seurat^71^ data objects to store, annotate and retrieve information from scRNA-seq experiments. We offer all functionalities in the R programming language whilst benefitting from the CellRank^31^ and deeptime^72^ python packages to carry out state transition and TPT inferences; sciCSR deploys these python packages at the backend using the R reticulate environment. Whilst some steps would benefit from the usage of high-performance computing (HPC) clusters (e.g. the enumeration of productive and sterile transcripts), all steps of the sciCSR pipeline can be performed on modern desktops and laptops. sciCSR allows users to perform CSR analyses on an “ad-hoc” basis just as other conventional downstream scRNA-seq data analyses and visualisation, avoiding the need to restart raw data processing from sequencing read FASTQ files to retrieve productive and sterile transcript counts as has been the case in previous analyses^11, 52^.

### sciCSR predicts CSR directionality in temporal scRNA-seq data

The design principle of sciCSR enables the use of a scRNA-seq “snapshot” of B cells to infer their CSR tendencies (i.e. the isotype[s] the cells are going to switch from/to) during immune response. We first tested this idea by utilising a published dataset by Kim et al.^73^ which profiled GC B cells following SARS-CoV-2 mRNA vaccination to monitor long-term B cell maturation. **Figure 4** shows data from two donors for which complete datasets with data from weeks 4, 7, 15 and 29 post first dose of vaccination are available, with the latter three timepoints sampling B cells from lymph nodes and profiling using scRNA-seq and scBCR-seq. We used sciCSR on week 7 scRNA-seq data of GC B cells and sought to predict the isotype distribution observed using week 15 scBCR-seq data, and similarly analysed week 15 scRNA-seq to predict week 29 scBCR-seq. The two donors displayed differences in their BCR isotype distributions (**Figure 4**b), and sciCSR successfully predicted these distributions using donor-specific GC B cells with high accuracy (median cosine similarity = 0.949) when comparing the TPT inward fluxes (i.e. amount of CSR towards each isotype) against the isotype distribution observed in scBCR-seq data at the subsequent timepoint (**Figure 4**c-d). The flux matrix and its associated p-values can be visualised in a bubble plot which breaks down the inferred CSR fluxes, revealing different switching sequences (IgG1 -> A1 -> G2 for donor 07, and IgG1 -> IgG4 accompanied by switching to IgA1 for donor 20) which can be validated with scBCR-seq data. sciCSR successfully predicts the directionality of CSR and uncovers nuances in CSR trajectories which are otherwise hidden in the scRNA-seq data space, given that these GC B cells do not appear transcriptomically distinct in the original analysis by Kim et al.^73^ (Supplementary Figure S4).

**Figure 4.**
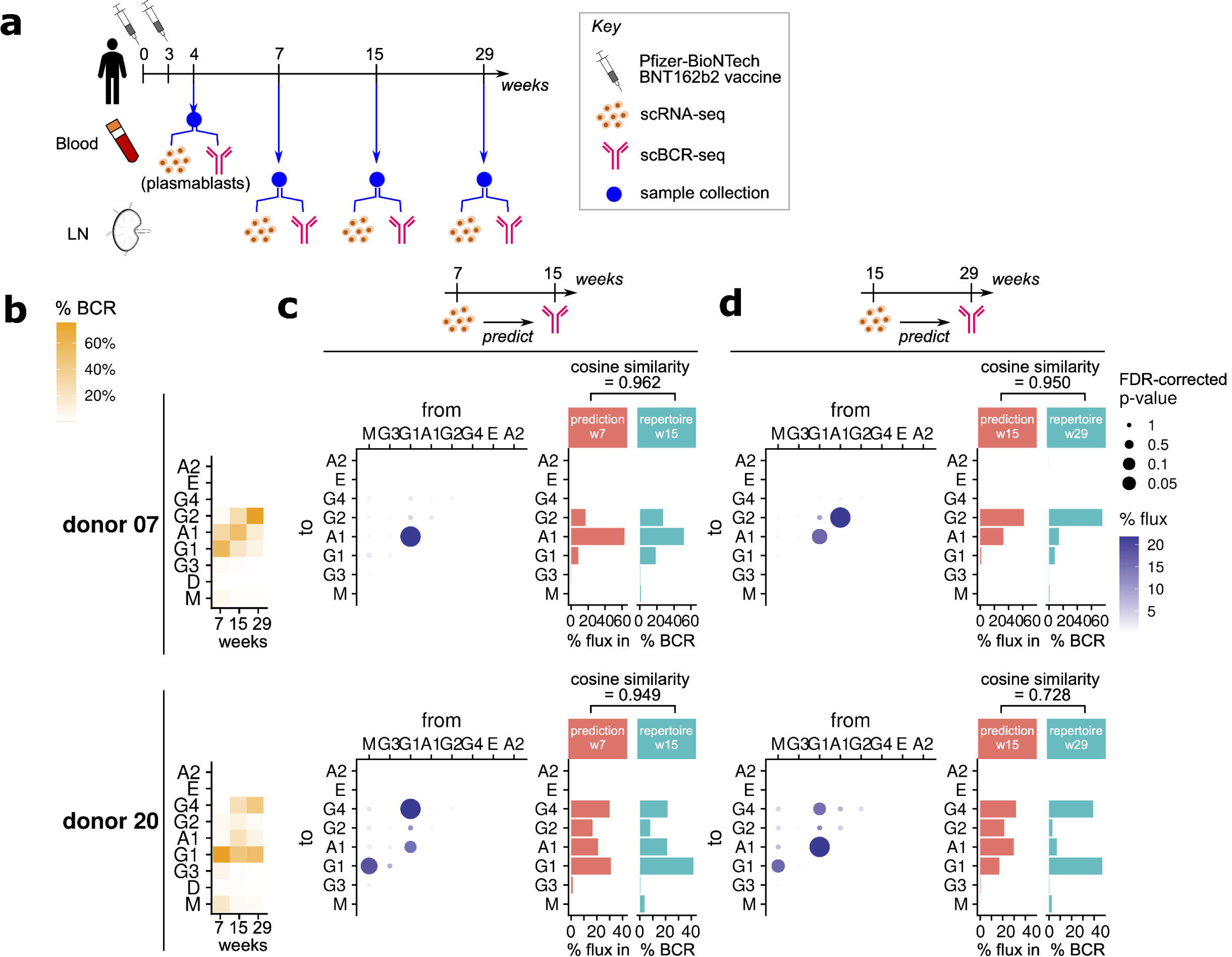
sciCSR predicts BCR isotype distribution in time-course immunisation data. (a) Schematic of immunisation and follow-up data collection from the Kim et al dataset. (b) Isotype distribution of GC B cells for donors 7 (top) and 20 (bottom), across weeks 7, 15 and 29 scBCR-seq data. (c, d) sciCSR-predicted CSR fluxes, based on Transition Path Theory (TPT), from and to eac**h** isotype. Predictions were made based on week 7 scRNA-seq data to predict scBCR-se**q** isotype distribution at week 15 (panel c), or using week 15 scRNA-seq data to predict week 29 scBCR-seq isotypes (d). *(left)* CSR fluxes expressed as a bubble plot. The colours of bubbles correspond to the amount of flux between the given pair of isotypes predicted by sciCSR, and their sizes correspond to statistical significance of these fluxe**s** compared to predictions generated from randomised transitions (see main text and Materials and Methods). *(right)* The sciCSR-predicted switching events to each isotype, represent by total inward TPT fluxes towards each isotype, is compared against the observed BCR isotype distribution at the subsequent timepoint. These two distributions were compared using cosine similarity.

### sciCSR demonstrates differences in CSR introduced by perturbation experiments

We reason that sciCSR can be an ideal tool to analyse functional genomics experiments which aim to uncover gene effects on B cell maturation by introducing perturbation to the system. To investigate this in greater detail, we first collected from the literature scRNA-seq data of gene knockouts with reports of CSR effects. **Figure 5** shows application of sciCSR to analyse three such datasets, on mice with three genes *(Aicda* ^74^, *Hspa13* ^75^ and *Il23* ^76^) knocked out, either at a whole-organism level or conditionally in Cd19+ B cells. In the original reports, *Hspa13* and *Aicda* knockouts decreased both CSR and SHM^74, 75^, whilst *Il23* knockouts biased the cells away from IgG2b^76^. Applying sciCSR on the knockout and wild-type mice scRNA-seq data, we recapitulated the outcomes reported in these studies (**Figure 5**a-c). Since the TPT workflow implemented in sciCSR accepts any one of RNA velocity, CSR or SHM ordering of cells as inputs, we asked whether these methods capture different information in the inference of cell state transitions. We considered the Hong et al. *Il23* knockout dataset^76^ where both scRNA-seq and scBCR-seq data were complete and available, inferred the cell-cell transition matrices using each method, simulated transition paths given the resultant Markov chains, and compared the frequencies these simulated paths visited each state in the data (**Figure 5**d). We observed that RNA velocity consistently provided different inference than to CSR and SHM, with the latter two methods capturing almost identical trajectories (**Figure 5**e), validating that sciCSR captures CSR information which is consistent with evidence from SHM in inferring transitions. Both represent B-cell-specific evidence to infer state transitions relevant to the system; given RNA velocity methods are known to give noisy inference in mature immunological cell types^24, 32^, sciCSR could serve as a viable alternative by harnessing CSR and SHM to analyse B cell state transitions.

**Figure 5.**
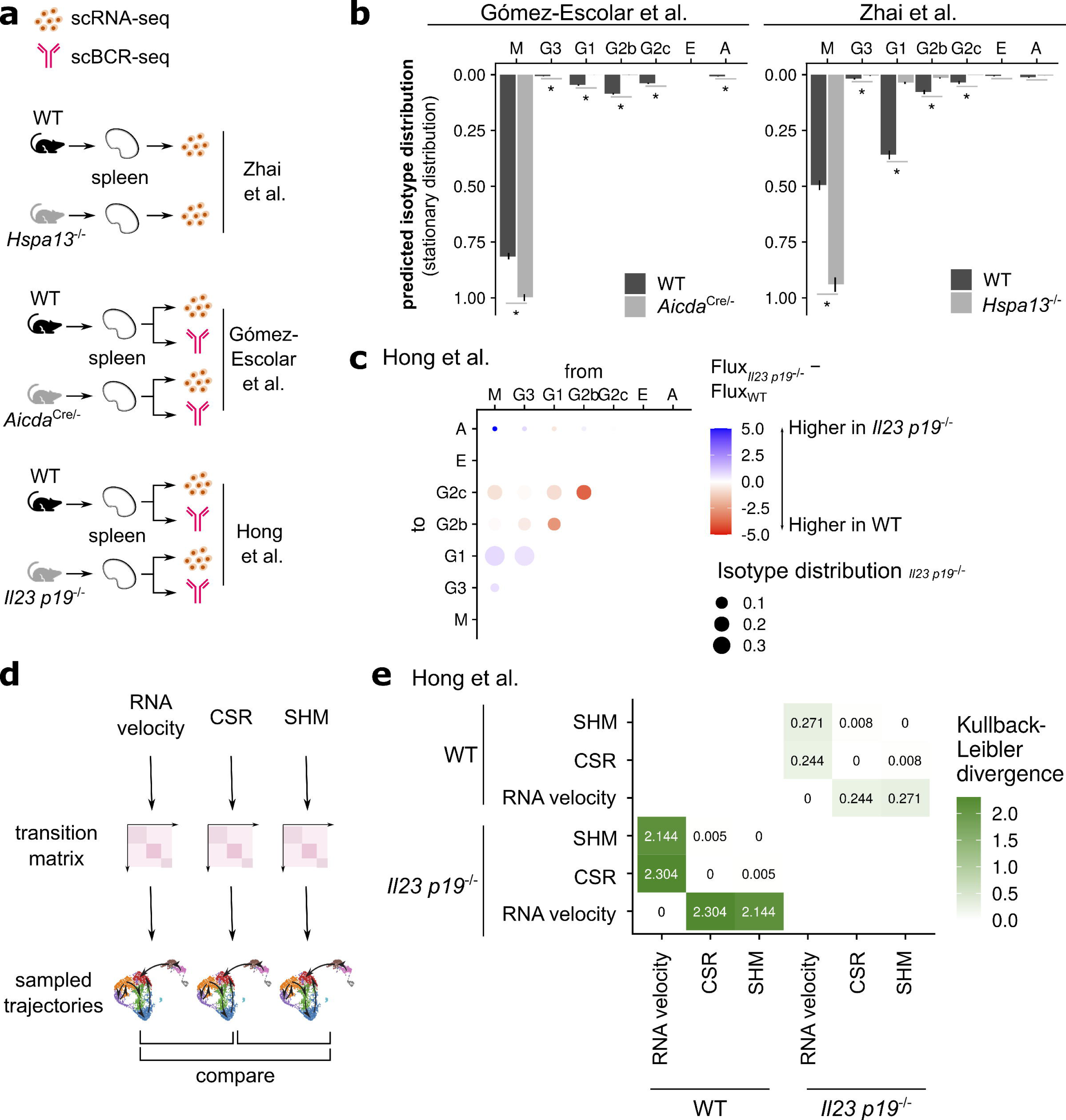
sciCSR recapitulates CSR effects in gene knockout scRNA-seq studies. (a) Schematic of scRNA-seq/scBCR-seq data from mouse gene knockout studies considered in this paper. Notations of knockout genotypes follow the respective publications. (b) Isotype distribution in the Gómez-Escolar et al. *(Aicda* knockout) and Zhai et al *(Hspa13* knockout) data. Stationary distributions predicted by sciCSR are proxies of isotype distribution at equilibrium. sciCSR were applied separately for wild-type and knockout cell**s.** Error bars represent 95% confidence intervals obtained by bootstrapped sampling of cells for each isotype. *, false-discovery rate (FDR) adjusted p-value < 0.05 for comparison of wild-type and knockout cells proportions for each isotype, evaluated using two-sample Z-test for proportions. (c) Difference in sciCSR-predicted CSR fluxes in the Hong et al. data, comparing the *II23 p19-’-* and wild-type (WT) cell subsets of the scRNA-seq data. Isotype combinations with higher levels of CSR in the WT cells are highlighted in red. Bubble sizes are scaled by the isotype distribution in the *Il23 p19^-^’^-^* data. (d) Illustration of comparing transitions inferred using different biological information. Transition matrices were constructed using CSR, SHM, and RNA velocity (calculated using scVelo) information. The resultant Markov chains derived from each transition matrix were used to sample trajectories of cell state transitions; the frequencies of visiting each cell state in these sampled trajectories could be directly compared. (e) Comparison of transitions inferred based on the schematic in panel (d), on the Hong et al. *Il23 p19* knockout dataset using RNA velocity, CSR and SHM information separately on the WT and *Il23 p19-’-* cells. The distributions of frequencies of visiting each cell cluster derived from the sampled trajectories are compared using Kullback-Leibler divergence.

## Discussion

A lack of accurate tools to study cell state transitions in mature cell types, specifically immune cells, preclude computational analyses of these processes in scRNA-seq datasets of important biological phenomena such as immune response to vaccines, pathogens or malignant cells. sciCSR addresses this gap by implementing a novel approach for studying B cell state transitions. The uniqueness of our approach focuses on extracting B cell-specific information (expression of sterile/productive IgH transcripts, somatic hypermutation level) from scRNA-seq/scBCR-seq datasets, whilst benefitting from the well-documented and comprehensive software package CellRank^31^ to provide underlying implementations which infer cellular trajectories. We introduce TPT^70^ to analyse Cell Rank-inferred transition matrices. Inspired by analysis of molecular dynamics simulation data, TPT can act on the transition matrices inferred using B cell-specific information, to describe the collective CSR dynamics of B cells. TPT and the generation of null transitions matrices (where transition probabilities are randomly reshuffled) offer a principled way to assess the robustness of the inferred transitions. sciCSR is robust in recovering transitions between small populations (of size as small as groups of 10 cells, Supplementary Note 1), making it ideal to study transitions in rare B cell populations (e.g. antigen-specific B cells from peripheral blood).

We believe a standout feature of sciCSR is its design principle of utilising cell-type-specific biological features to inform trajectory inference: many commonly used tools for inferring cellular developmental trajectories are agnostic to the biological system of study. CellRank by default uses RNA velocity to build cell-to-cell transition matrices; here we derived a custom cell ordering based on B cell biology and used this as input to CellRank to infer transitions. RNA velocity information was known to be sparse in mature immune cell types in the PBMC population^24, 32^; whilst RNA velocity has previously been applied to B cell scRNA-seq datasets^10, 11^, its application is noticeably less widespread compared to systems in developmental biology, potentially explained by the noisy inference of velocity models in many mature immune cell populations^24, 32^. Tools which utilise system-specific biological knowledge to learn vector fields which describe cellular dynamics have recently received more attention, owing to their potential in overcoming the limitations of general trajectory inference tools for specific biological systems. Some necessitates collection of orthogonal data types, e.g. barcode-based lineage tracing of the same cell population sampled using scRNA-seq (PhyloVelo^77^); here, whilst B cells represent an attractive system to apply such methodologies (due to the fact that these phylogenies can be readily reconstructed from scBCR-seq V(D)J sequences), the limited number of B cells represented in many currently available scRNA-seq libraries implies restricted sampling of these lineages, which may pose challenges to the subsequent trajectory inference step. One approach (Pseudocell Tracer^78^) considered the BCR isotype expression profiles and trained deep-learning-based generative models to overcome the sparsity of cell states in the data. These generative models were then used to map out the inherent CSR trajectory in the generated latent space. This represents a viable approach to model CSR in scRNA-seq data; however, it lacks a publicly available implementation which can be readily used in analysing new datasets. Here we build upon CellRank^31^ and interface with well-supported scRNA-seq data analysis packages in R and python (Seurat^71^, scanpy^79^, etc.) to provide a method which also incorporates the underlying B cell biology based CSR and SHM signals from the data. We have evaluated the performance of sciCSR in a range of scRNA-seq datasets generated under common model systems used in immunology such as vaccination and mouse knockout models. With these datasets we show that sciCSR offers accurate predictions of CSR which can be verified by measuring the BCR isotype expression subsequent to perturbations.

The major innovative feature in sciCSR is the enumeration of productive and sterile IgH transcription in single cells, which can be applied in any human and mouse B cell scRNA-seq dataset. Whilst analysis of sterile transcription was previously attempted in some B cell scRNA-seq studies, they were performed at the stage of raw data pre-processing^11, 52^, without the relevant code made available in the associated publications. sciCSR allows this analysis to be performed on an ad-hoc, *a posteriori* basis as for other common scRNA-seq data analysis tasks. Similar to count data for many other transcripts in the transcriptome, counts for productive and sterile IgH transcripts are sparse, especially for the productive transcripts (Supplementary Figure S3); this can be mediated with parallel scBCR-seq libraries obtained for the same cells, which are increasingly popular in studies focussed on B cells, particularly their antigen-binding features^11, 73^. The deconvolution into sterile and productive transcripts is useful in highlighting the diversity of CSR tendencies between B cell subsets (for example, depending on B cell subsets, up to 25-30% of cells express sterile transcripts of more than 1 isotype in human B cell scRNA-seq atlases we have analysed, see Supplementary Figure S3).

A recent analysis by Horton et al. ^50^ uses lineage tracing to study the establishment of B cell fates, and found that the CSR fate of a B cell is largely independent from its predecessors; CSR fate is instead determined by a combination of AID expression and sterile transcription^50^, as previously noted by others^35, 39, 51^. In agreement with this, we observed that the presence of sterile transcripts within the same BCR clonotype is a noticeably weaker indicator of BCR isotype, in comparison to the control scenarios of recalling the BCR isotype of a given cell (as captured in repertoire sequencing data) using productive and sterile transcripts from scRNA-seq data of the same cell (Supplementary Figure S5). We believe that the enumeration of productive/sterile transcripts opens the door to further explore the basic mechanisms of CSR: the enumeration protocol in sciCSR can support the use of scRNA-seq to study the variegation of CSR fates, and motivate mathematical modelling approaches to quantify CSR likelihoods beyond the analysis presented in Horton et al^50^; coupling with experimental approaches to study single-cell epigenetic landscape of B cells^80^ and computational approaches to infer gene regulatory activities^81^, the enumeration of productive and sterile transcripts could contribute towards understanding how sterile transcription of different isotypes are differentially regulated, how this reconciles with their observed baseline expression levels, and how different molecular stimuli could modulate sterile transcription and ultimately fine-tune CSR *in vitro* and *in vivo*.

sciCSR is a freely available R package providing implementations to build inference of CSR and B cell maturation using scRNA-seq data; all functionalities can be performed at the R frontend to provide users, particularly those accustomed to scRNA-seq data analysis in R, with a unified data analysis pipeline whilst benefitting from trajectory inference tools written in python, such as TPT and the implementations offered by CellRank^31^. The enumeration of productive and sterile IgH transcripts does have its technological limitations: it necessitates usage of the 5’-biased read sampling protocol which would currently preclude datasets generated using the more common 3’ protocol and the use of spatial transcriptomics data. More detailed analyses in the future (e.g. identification of latent features of CSR states which are also detectable using 3’ data) will extend the utility of sciCSR to different technological platforms. We believe sciCSR offers a starting point to model B cell maturation in scRNA-seq data; these models can be further analysed to understand the molecular cues of CSR and different steps of the maturation process, their regulation *in situ* within tissues, and their dysregulation in diseases.

## Data availability

All datasets used in this work are publicly available: Kim et al.^73^ (Gene Expression Omnibus [GEO] entry GSE195673), Gómez-Escolar et al.^74^ (GSE189775), Hong et al.^76^ (GSE145922), Zhai et al.^75^ (ArrayExpress accession E-MTAB-8280), Stewart et al.^10^ (E-MTAB-9544), King et al.^11^ (E-MTAB-9005), Mathew et al.^13^ (E-MTAB-9478 and E-MTAB-9491) and Luo et al.^14^ (E-MTAB-10081). For GEO entries raw FASTQ files were downloaded from the associated Sequence Read Archive (SRA) entries.

## Code availability

Source code for the sciCSR package is available at: https://github.com/Fraternalilab/sciCSR. Documentation and vignettes can be found in the github repository.

## Acknowledgement

This work was funded by the Biotechnology and Biological Sciences Research Council (BB/T002212/1) The funders had no role in the collection and analysis of the samples, in the interpretation of data, in writing the report, nor in the decision to submit the paper for publication. The authors would also like to acknowledge Dr. Jens Kleinjung for his critical reading of the manuscript.

## Materials and Methods

### The sciCSR algorithm

sciCSR is a R package to annotate productive and sterile IgH transcripts given scRNA-seq aligned reads, and process scRNA-seq (and scBCR-seq if available) data to infer state transitions using CSR and SHM information. The inputs to sciCSR are (1) BAM files of the processed and aligned scRNA-seq reads; (2) processed and normalised cell-by-gene count matrix corresponding to the BAM files, and (3) optionally, associated scBCR-seq data for the same set of cells. If (1) and (2) are available, sciCSR will extract CSR information and use this to infer cell state transition. If (3) is also provided, sciCSR will extract both CSR and SHM information and users can opt to use either in the inference. For notational convention let *N* be the number of cells and *G* be the number of genes transcriptome-wide recorded for a given scRNA-seq dataset. sciCSR aims to:

(i) Enumerate productive and sterile immunoglobulin heavy-chain (IgH) transcripts, as defined in the main text and illustrated in Figure 1c, for each cell. These results produce a separate cell-by-transcript count matrix where for each IgH isotype, two transcripts will be listed, one corresponding to the productive transcript of the given isotype, and another corresponding to sterile transcripts. For a species with *S* isotypes, this will yield a 2S × *N* matrix which we will refer to as “isotype matrix” in the following text.

(ii) Use the isotype matrix or, if scBCR-seq data is available, the SHM level as input to generate a cell ordering, as a “pseudotime” input to CellRank to infer a cell-by-cell transition matrix. CellRank^31^ is used in accordance with author-recommended protocols, i.e. the final cell transition matrix combines the matrix inferred using the *k-* nearest neighbour (kNN) network that describes transcriptomic similarity between cells, and the matrix inferred using CSR/SHM information. CSR/SHM information is used as bias to impose directionality on the kNN-based transition matrix.

(iii) Infer the fluxes between either cell clusters or isotypes by applying Transition Path Theory (TPT) on the transition matrix.

### Mapping positions of sterile transcription at the IgH genomic loci

The problem of identifying sterile IgH transcripts given scRNA-seq reads mapped to the reference genome is non-trivial, since sterile transcripts are not annotated in standard genome references such as those provided by Ensembl or UCSC genome browser. The distinction between productive and sterile IgH transcripts lies in the transcription start site (TSS): productive IgH transcripts are transcribed beginning at positions 5’ to the Ig variable (V) gene in the leader region, whilst sterile transcripts begin at some position 5’ to the first constant region (C) exon^42,46–48,82^. We first clarified the exact genomic position of TSS for the sterile transcript for every human and mouse IgH C gene with the exception of IgD where sterile transcription is not well understood. We collected from the literature a set of DNA genomic sequences isolated using traditional molecular cloning and restriction enzyme analysis, of the promoter region of sterile transcripts observed in experimental and/or clinical models of csR^46–48,83–89^. These sequences are mapped to the hg38 (for human) /mm10 (for mouse) reference genomes using BLAT^90^ (v.36x4). Supplementary Table 1 lists the positions of sterile transcript TSS; this table is stored in the sciCSR package to guide the extraction of aligned scRNA-seq reads for enumerating productive and sterile transcripts (see below).

### Re-analysing scRNA-seq read alignments to distinguish productive and sterile IgH transcripts

The first step of sciCSR reads in user-supplied BAM files where aligned reads are listed with additional fields noting their associated cell (with tag “CB”) and molecular (tag “UB”) nucleotide barcode; these tags which are used in 10X cellranger-generated output BAM files are set as default. The BAM file is scanned three times, each time extracting reads mapped to a different genomic location: (1) the exonic regions encoding any IgH V, diversity (D) or joining (J) gene; (2) the exonic regions encoding any IgH C gene, and; (3) the 5’ region starting from the sterile transcript TSS to the site immediately preceding the start site of the first C exon (hereafter “5’ C”). In each scan sciCSR extracts the aligned genomic positions for reads fulfilling each criterion listed above. Spliced reads which merely span across a given genomic range without actual matches to the genomic region of interest, as well as those without cell and/or molecular barcodes, are removed in each scan. We next implemented the following scheme to summarise these three lists of aligned reads to discern, for each molecule with a unique combination of cell and molecular barcodes, whether they represent productive or sterile IgH transcripts (Figure 1c):

(i) A “productive” transcript must have at least 2 reads, one mapping to the VDJ genes and another to the C exonic regions;

(ii) A “sterile” transcript must have at least 1 read mapping to the 5’ C region, and optionally to the C exonic regions, but no read mapping to the VDJ genes, and;

(iii) Transcripts where insufficient information exists to be classified as productive or sterile are labelled “uninformative”.

The number of cell-molecule barcode combinations were enumerated for each productive/sterile/uninformative IgH transcript type. Discounting the uninformative transcripts, this gives, for each cell, a gene count vector of length *2S* where *S* is the total number of IgH C genes for the given species. Concatenating these vectors column-wise yield a *2S×N* matrix where *N* is the total number of cells. We term this matrix the “isotype matrix”.

### Validating workflow to identify productive and sterile IgH transcripts

We validated the enumeration of sterile and productive transcripts by utilising a simulated dataset where we sampled sequencing reads from the human IgH genomic locus. Comparison between the ground-truth distribution of reads we sampled and the isotype distribution reconstituted after aligning these reads to the reference genome would indicate whether the workflow can distinguish between productive and sterile IgH transcripts. To generate this simulated dataset we concatenated the DNA sequence for the VDJ region of the antibody CR3022 (GenBank accession DQ168569)^91^ with the exonic regions of each human IgH C gene (GRCh38 reference genome from Ensembl); this set of sequences would be the full-length productive transcripts. For sterile transcripts, the genomic sequences (genome build hg38) of each human isotype between the sterile transcript TSS and the end of the C gene coding region was retrieved from the UCSC genome browser application programming interface (API). All productive/sterile transcript sequence constructs are included in Supplementary Data 1. We next designed scenarios where cells express either one specific sterile/productive transcript, or a combination of sterile/productive transcripts of different isotypes at given proportions; all such scenarios were shown in Figures 2b and 2c. The R package polyester^92^ (v1.29.1.1) was used to sample reads based on these ground-truth sampling proportions, using the function “simulate_experiment_countmat”. To mimic experimental data where reads are biased towards either the 3’ or the 5’ ends of transcripts, we included the positional bias model from Li and Jiang^93^ as an argument for polyester to generate sets of reads biased to the 3’ end, and the mirror image of the Li and Jiang model as input to generate 5’-biased reads. All other parameters to run polyester were unchanged. A total of 1 million reads was sampled for each designed scenario. The sampled reads were subjected to alignment to only the chromosome 14 genomic sequence from the GRCh38 reference genome, using either HISAT2^67^ (v2.2.1) with default parameters, or STAR^66^ (v2.5.1.b) using the following parameters: “--outSAMmultNmax - 1 --readNameSeparator space --outSAMunmapped Within KeepPairs”. The resultant BAM files were then subject to productive/sterile transcript enumeration using the pipeline described above. We attempted in parallel quantification using a pseudoalignment tool for validation; salmon^94^ (v1.9.0) was used by supplying a custom genome containing the GRCh38 chromosome 14 genomic sequence and all reference human IgH productive/sterile transcript set described above. We found that quantification by salmon was noticeably more inaccurate compared to those based on applying our pipeline to HISAT2/STAR-generated alignments. They were therefore not included in the results shown in Figure 2.

### Combining scRNA-seq and scBCR-seq data

sciCSR provides functionalities to merge scRNA-seq and scBCR-seq measurements obtained in parallel for the same set of cells using the 10X Genomics technology. sciCSR expects the 10X scRNA-seq and scBCR-seq data be pre­processed using the 10X cellranger software suite, and the resultant scRNA-seq gene count matrix be imported in R and stored as a Seurat data object. We added, as separate columns in the metadata slot of the Seurat data object, sequence annotations in the filtered contigs comma-separated value (CSV) files (i.e. those with filenames “filtered_contig_annotations.csv”) from the “cellranger vdj” run. The filtered contig annotations were cleaned for cases where more than one heavy and/or light chain sequence was associated to the same cell, taking only the sequence with maximum unique molecular identifier (UMI) count as representative. This data table is organised such that sequences are organised into one cell per row, listing separate heavy and light chain annotations as columns. By default the following annotations are retained and merged into the Seurat object metadata: V, D, J, C gene annotations, binary variables of whether sequences are productive and full-length, the CDR3 nucleotide and amino acid sequences, and the total read count and UMI count associated with each heavy/light chain sequence.

### Deriving CSR potential

For CSR, sciCSR makes use of the productive/sterile isotype matrix to derive a score which ranks cells from naïve to memory/terminally differentiated. The intuition is that the isotype matrix can be decomposed into “signatures” of isotype expression which are representative of naïve/memory cell types; these signatures can then be used to deconvolute the isotype matrix and score the resemblance of each cell to naïve or memory state, in terms of their IgH transcription profile. To guide the definition of these isotype signatures we first applied the sciCSR productive/sterile transcript quantification to scRNA-seq data of B cells at reference B cell states; these reference B cell atlases were compiled for human and mouse (see heading *“Reference atlases”* under “Datasets” in this Materials and Methods section). These datasets were chosen as reference B cell atlases to cover a wide range of B cell states found in circulation and in secondary lymphoid organs, and, wherever possible, sampled from healthy subjects. We next applied non-negative matrix factorisation^68^ (NMF) separately on the human and mouse data, to decompose the isotype matrix *T* (of dimension *2S* (total number of isotypes) × *N* (number of cells)) into two matrices:

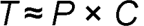

The matrix *P* is of dimension 2S × *H,* where *H* is the number of isotype signatures. *C* is a matrix of dimension *H×N.* Each isotype signature consists of weights indicating the importance of the given productive/sterile transcript for a cell type of interest, whilst an entry of the *C* matrix *Cĳ* would refer to the contribution of isotype signature *i* in cell *j.* Note the approximation sign in the equation; applying NMF is to jointly estimate *P* and *C* which together approximates best the input matrix *T.* Our goal here is to obtain one signature (i.e. column) *v* from the matrix *P* which either biases towards memory B cells and/or terminally differentiated cells in the lineage such as plasma cells (i.e. c_v;_· increases as cells go from naïve to memory), or one that is biased to the sterile/productive transcripts of IgM (i.e. the developmental trajectory would go inversely of this signature) to rank the cells. In principle we could obtain signatures specific to IgG/A/E using this method; here the IgM-biased signature was deemed preferable as it could be applied to different biological contexts where any switching events away from IgM were of interest.

We first compared the signature matrix *P* obtained using NMF upon incrementing the value the number of signatures *H* from 2 to *S*, and observed that as *H* increased, we obtained more fine­grained signatures eventually ending up with signatures composed of only 1 isotype (Supplementary Figure S6). For the human reference atlas setting *H* = 2 yields a naïve B cell signature biased towards IgM, and a memory signature which was a combination mainly of different IgG subtype productive/sterile transcripts (Supplementary Figure S6). For the mouse atlas setting *H* = 2 did not yield separate IgM+ and IgM-signatures, but for *H* = 3 we obtained an IgM_sterile_+, an IgM_produ_ct_i_v_e_+, and an IgG-biased signature (Supplementary Figure S6). We therefore directly took the human *P_H_* = _2_ matrix as the reference human signature matrix, and for mouse we sum over the IgM_sterile_+ and IgMp_roductive_+ signature weights to generate a two-signature (IgM-biased, IgG-biased) *P* matrix as the reference mouse signature matrix. These *P* matrices are stored to score cells from a new, user-supplied, dataset *T’,* using non-negative least square (NNLS) regression, i.e. estimating *T’∼P× C’*using *P* trained on the reference dataset *T.* The entries *c’_vj_* in the *C’* matrix corresponds to the contribution of the naïve (i.e. IgM-biased) signature *v*, trained on the reference atlas, in cell *j* in the user-supplied data. cýyfor all *j* were taken to derive the CSR potential by scaling into the range [0,1] and reversing the scale such that switched cells (i.e. less naïve-like) have a higher CSR potential:

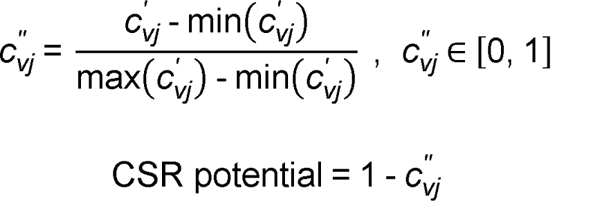

NMF was performed using the R NMF package (v0.24.0) using the “nmf” function, applying the alternating least square approach from Kim and Park^95^ (argument “methods = ‘snmf/r’”). The “nmf” function attempted 10 NMF fits and the best (i.e. the smallest approximation error) fit was taken as the final signature matrix to be stored and used to score user-supplied datasets. For the human atlas, all cells were considered and the “nmf” function was applied once; for the mouse atlas, to reduce memory usage, the “nmf” function was called 10 times, each time using a different, randomly selected set of 20,000 cells from the atlas to fit 10 NMF. Signature matrices from the best NMF fit from each of the 10 subsetted datasets were summarised as element-wise mean values to derive the final signature matrix for storage and decomposition of user-supplied productive/sterile transcript counts. NNLS was performed using the R package nnls (v1.4).

### Deriving SHM level

If scBCR-seq data has been integrated with scRNA-seq results, sciCSR calculates the SHM level of each cell as follows:

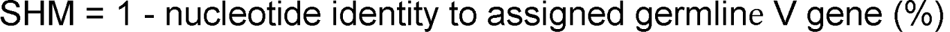

This value is subsequently scaled into range [0,1] similar to the derivation of the CSR potential.

### Applying CellRank

The CSR and SHM ranking of cells were taken as “pseudotime” orderings, to generate cell-cell transition matrix using the CellRank^31^ (v1.5.1) “PseudotimeKernel”. We followed the CellRank recommended protocols and generated a final transition matrix *M* for each dataset by combining transition matrices using two different sources of information:

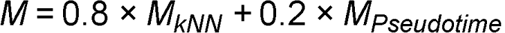

where *M_kNN_* is the transition matrix (“ConnectivityKernel” in CellRank) derived by considering the kNN network calculated using scanpy (v1.9.1), and *Mp_seudotime_* is the transition matrix derived from using the CSR/SHM information as pseudotime ordering of cells. The respective weights of *M_kNN_* and *Mp_seudotime_* were defaults for combining kernels in CellRank^31^.

### Transition Path Theory (TPT)

TPT is a statistical framework to analyse the properties of transition paths governed by a discrete-state Markov chain^56, 69, 70^. The major premise of TPT is, since transition paths between states can adopt a variety of behaviours (e.g. to reach state *B* from *A,* this can be accomplished via a direct jump, or a more convoluted path involving some other intermediate state(s), or simply fail to reach *B* and return back to *A*), it is important to describe the statistical properties of an ensemble of transition paths, from a source state to a target state (both defined by the user), in order to capture more truthfully the transition dynamics in the system^69, 70^. TPT is a well-established technique in physics and chemistry to study dynamics of chemical reactions and conformational changes of molecules^56, 70, 96^; the latter has been adopted in computational biology in the analysis of molecular dynamics (MD) simulation trajectories^53–55^. We used the python package deeptime^72^ (v0.4.2) (which powered the backend calculations of TPT in the MD analysis package PyEMMA2^53^) for TPT calculations. deeptime implements TPT in a python class “ReactiveFlux” to fit TPT on the CellRank-derived transition matrix *M*, such that it describes the transition dynamics of the cells from a source to a target state indicated by the user. These source/target states can either be cell states defined by the cell cluster labels users typically give to scRNA-seq datasets, or BCR isotypes specifically for the analysis of CSR dynamics, or any arbitrary grouping defined by the user. Since the transition matrix *M* from CellRank has dimension *N × N* where *N* is the total number of cells, a grouping (“coarse-graining” in the deeptime package) step is required to group fluxes such that they can be interpreted between clusters of cells. We fit TPT on the *N× N* transition matrix giving vectors of cell indices corresponding to the source and the target state respectively, and then use the ReactiveFlux.coarse_grain() function in deeptime to coarse-grain the fluxes at the cell cluster level. These “clusters” are supplied by the user; they can either be cell identities defined by clustering the transcriptomic data, or BCR isotypes.

The ReactiveFlux object computes various quantities typical of Markov chain and TPT analysis, but we deem the following the most informative about cell state transitions:

(1) Gross flux *F*, a *n× n* matrix where *n* is the total number of cell states. Each element *fĳ* describes the amount of flux that flows from state *i* to state *j.* Importantly, TPT provides flux estimates for any pair of states, not just limited to direct fluxes between the source and target states. We adopted the recommendations in PyEMMA2^53^ to interpret these as relative fluxes, i.e. instead of the absolute flux values these are normalised as 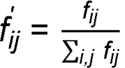 which describe the proportion of total flux in the system which flows from state *i* to *j.* These fluxes are the indicators of transitions and can be visualised with e.g. projection of arrows on dimensionality reduction plots, typical in scRNA-seq transition dynamics analyses such as RNA velocity^19, 20^.

(2) Stationary distribution *π.* This is a vector of length *n*, where each element is a probability describing the likelihood to reach a given state when the system reaches equilibrium over a long period of time. Intuitively the stationary distribution is informative in the comparison of transition dynamics observed in perturbation experiments where the perturbation (e.g. a gene knockout) yields a discernible effect to the likelihood to sample certain states. All elements in *π* sum to 1, i.e. *Σ_n_π = î*.

(3) Mean first passage time (MFPT) between the source and target states. The MFPT is the inverse of the transition rate between the source and target states and quantifies the efficiency of transition given the data.

(4) A list of transition pathways with intermediate states from the source to the target, with probabilities of their traversal in order to reach the target from the source.

These results are provided when users execute the fit_TPT() function in the sciCSR R package together with analogous null/bootstrapped estimates to assess statistical robustness (see below).

### Interpreting TPT

TPT fluxes are descriptions of transitions between states estimated by considering the frequencies each state is sampled in the data and the transition behaviours of each cell as specified in the input transition matrix. Since the goal of TPT is to evaluate possible pathways to traverse the state landscape to reach a user-specified target state from a source state, the choice of source and target states influence the TPT inference results (Supplementary Figure S8). Specifically, the magnitude of flux tends to be higher for fluxes involving either the chosen source or target states, since the algorithm is designed to consider pathways that necessarily pass through these states as required by user definition. It is therefore important to choose appropriate states with the aim of describing the fluxes and stationary distributions of all states in the system. Here we chose source and target states which were (1) sampled in the data and (2) as close to the ultimate start/end points of the relevant biological process as possible. For example, for analysing CSR we would want to set the source state as IgM and the target state as IgA2 (human)/IgA (mouse) given these isotypes were observed in the data; transition paths involving any possible pairs of isotypes in the middle of the locus would then be traversed, some of which would be of interest to the analysis. For analysing transitions between cell clusters, we chose the cell cluster representing naïve B cells (if available) as the source state, and the plasma cell/plasmablast cluster as the target state. We reasoned that in most use cases users would have an intuition of the observed states and be able to choose source and target states using prior knowledge.

### Pruning TPT results

While quantities listed in the section “Transition Path Theory (TPT)” are typically analysed in usage of TPT in biological settings (e.g. in MD analysis^53, 54^), we reasoned that additional measures were necessary to prune the TPT results and give uncertainty estimates to fluxes and stationary distribution quantities to aid interpretation. In applying TPT to infer CSR, backward fluxes (that is, flow from a given isotype to another which is 5’ to itself) are excluded as these transitions are physically prevented via genomic DNA excision during CSR. To evaluate the statistical significance of fluxes, we estimated a p-value for each entry in the gross flux matrix *F* by one-way comparison of whether the observed flux is greater than those obtained by randomly reshuffling the input transition matrix to TPT. We reshuffled columns of the transition matrix for *t* times (by default *t* = 100; users can change this number as appropriate) and fit *t* TPT models. The randomised fluxes were modelled as a Gaussian distribution and a one-way *z*-test was performed to derive p-values for each element of the gross flux matrix. The resultant p-values, after multiple-testing correction implemented using the FDR method with the p.adjust function in R, give confidence on how likely the observed fluxes are due to the structure of the transition matrix rather than merely the distribution of cell-types/isotypes in the system. For the stationary distribution, we performed boot-strap sampling of cells for each cluster label and summed over their individual stationary distributions, to obtain cluster-wise bootstrapped samples of stationary distributions.

### Visualising cell state transition inference results

sciCSR offers the following visualisation tools to present the TPT results: first, sciCSR provides function to depict the stationary distribution of states as a bar plot, with the 95% bootstrapped confidence intervals as error-bars. Second, for fluxes, users can project the inferred transitions onto either a grid of arrows or velocity streams, akin to RNA velocity plots generated using velocyto or scVelo. sciCSR uses the scVelo plotting facility in python in the backend and outputs the figure in R console. This visualisation is applicable for TPT inference where the cluster labels are used to group the cells. For analyses where cells groupings are not distinct on the dimensionality reduced space, users can also visualise fluxes as a bubble plot where each bubble corresponds to transitions between any pair of states; the bubble colour corresponds to the amount of flux, and bubble size indicates statistical significance (given by - log (adjusted p-value)). This visualisation is applicable to CSR-specific analyses grouping cells by BCR isotypes.

### Comparing transitions inferred using different biological information

Since sciCSR allows users to supply either CSR or SHM pseudotime orderings to infer cell transitions, and the underlying CellRank algorithm supports transition inference using RNA velocity information^31^, we reason that the CellRank transition matrices derived using these difference sources of information can be directly compared. To do so we set up separate Markov chains using the transition matrices, sample random walks from these Markov chains, and compared the frequencies of visiting each cell state in the sampled walks to ascertain whether these different transition matrices capture similar information regarding cell state transitions. For each transition matrix, a “markovchain” object (using the R package markovchain^97^, v0.9.0) was created and a total of 1,000 walks (users can change this default parameter) were sampled using the “markovchainSequence” function. The resultant frequency matrices which record the number of times each state is visited in the random walks are compared using either Kullback-Leibler divergence or Jensen-Shannon divergence, both calculated with default parameters using the R philentropy^98^ package (v0.5.0).

### Implementation

The sciCSR pipeline has been implemented as a R package. scRNA-seq data are handled using data objects defined in the Seurat (v4.0.6) R package^71^; sciCSR directly extracts and modifies data stored in the Seurat data objects, including inserting the isotype matrix as a separate “assay” in the Seurat object, and adding and modifying columns of the metadata. The inferences of state transitions using CellRank^31^ (v1.5.1) and TPT are implemented in python (v3.9.12) which sciCSR directly deploys under the R reticulate environment; sciCSR calls the R package SeuratDisk (v0.0.0.9020) (https://github.com/mojaveazure/seurat-disk) to convert between R Seurat object and python AnnData files as input to CellRank. TPT is fitted to the CellRank transition matrix using the python package deeptime^72^ (v0.4.2). A function within the R package (“prepare_sciCSR”) enable users to set up a working conda environment containing these python packages; the RNA velocity python package scVelo^20^ (v0.2.4) is also included in the environment to allow users to run CellRank using RNA velocity information for direct comparisons against the transitions inferred using CSR/SHM information as implemented in sciCSR.

### Datasets

#### Data preprocessing

Unless otherwise stated, the datasets below were processed as follows: raw FASTQ files were aligned to either the GRCh38 (for human data) or mm10 (for mouse) reference genomes and cell-by-gene count matrices were generated using 10X Genomics cellranger (v7.0.0). For datasets with scRNA-seq and scBCR-seq data available in parallel, the “cellranger multi” option was used to simultaneously call cells with both transcriptomic and VDJ data; otherwise the “cellranger count” option was used. Reference genome annotation (version 3.0.0 for both human and mouse) was downloaded from the 10X Genomics cellranger website. The “filtered_feature_bc_matrix” folder of cellranger output was read into R using Seurat::Read10X function as a cell-by-gene count matrix. Prior to generating a Seurat object which holds this count matrix, counts for genes encoding Ig V, D, J segments were summed for each cell as count for a meta-gene “Ig-vdj” to eliminate any effects caused by individual-/clonotype-specific VDJ gene usage to downstream clustering and differential expression analyses. The count matrix was then subsetted such that only features detected in at least 3 cells and cells with transcripts from at least 200 distinct features were retained. Dimensionality reduction and clustering analysis were performed stepwise as follows:

(1) removal of cells with the percentage of mitochondrial reads larger than 10%;

(2) log-normalisation using Seurat::NormalizeData with default parameters;

(3) identification of variably expressed genes using Seurat::FindVariableMarkers and removal of BCR-/TCR-specific genes from this variably expressed gene list as per Stewart et al.^10^;

(4) scale and center the normalised count data using Seurat: :ScaleData with default parameters;

(5) principal component analysis (PCA) using only the pruned variably expressed gene list;

(6) UMAP using Seurat::RunUMAP, calling the python umap-learn package and using correlation as the metric. We retained only the top *p* principal components, each of which explained at least 1.5% of the total variance;

(7) construct k-nearest neighbour (kNN) network using Seurat::FindNeighbors with default parameters and Louvain clustering on the kNN network. The resolution parameter is specific to each dataset and is indicated below separately.

#### Aicda gene knockout mice study from Gómez-Escolar et al^74^

This dataset contains scRNA-seq data from reporter mice where historical AID *(Aicda)* expression could be traced by sorting for expression of the fluorescent tdTomato (Tom) protein versus mice which are AID-deficient. Cells from the spleen were collected and prepared for scRNA-seq and scBCR-seq after immunisation with injection of the ovalbumin (OVA) protein. FASTQ files were downloaded from Sequence Reads Archive (SRA) accession SRP348368 and processed as detailed above, for sciCSR analysis of productive/sterile IgH transcripts. The transcript count matrix containing data for the other genes, as well as the cell metadata, were directly downloaded from the associated Gene Expression Omnibus (GEO) entry GSE189775 and imported using the Seurat package in R. sciCSR was applied to analyse transitions both between cell clusters and between BCR isotypes, separately for the AID-deficient and WT cells, using the signature-based CSR potential defined above. For inferring CSR, the following isotypes were chosen as the source/target states: WT - IgM (source), IgA (target); AID-deficient – IgM (source), IgG2b (target) (since for the AID-deficient cell subset no cells harbour isotypes further beyond IgG2b in the IgH locus).

#### Hspa13 gene knockout mice study from Zhai et al^75^

This dataset contains scRNA-seq data from splenocytes of *Cd19*-conditional *Hspa13* knockout *(Hspa13^-/^)* and wild-type mice. FASTQ files were downloaded from ArrayExpress accession E-MTAB-8280 and processed as detailed above. The dataset was filtered for non-zero expression of CD19 *(Cd19)* or CD20 *(Ms4a1)* to retain only B cells. A resolution parameter of 0.6 was used in Seurat::FindClusters to determine cell clusters. sciCSR was applied to analyse transitions both between cell clusters and between BCR isotypes, separately for the *Hspa13^-/^* and WT cells, using the signature-based CSR potential defined above. To infer CSR, IgM was chosen as the source and IgA as the target state in sciCSR.

#### Il23 gene knockout mice study from Hong et al^76^

This dataset contains scRNA-seq and scBCR-seq data from splenocytes of autoimmune BXD2 mice with knockout of the p19 component of *Il23* (hereafter *Il23 p19^-/-^)* and wild-type BXD2 mice. FASTQ files were downloaded from SRA accession SRP250728 and processed as detailed above. Unspliced and spliced transcripts were quantified using velocyto^19^ (v0.17.17), supplying both the mm10 General Transfer Format (GTF) file from cellranger references (see above) and the mm10 repeat mask GTF file obtained from the UCSC genome browser (RepeatMasker track) as arguments. RNA velocity estimation using the scVelo “dynamical” model was performed using the scVelo python package (v0.2.4) with default parameters. The filtered_contig_annotation files output from “cellranger vdj” were supplemented with the percentage identity to the germline V gene for each sequence by merging the cellranger filtered_contig_annotation files with the AIRR-formatted output of IMGT/HighV-Quest^99, 100^ (accessed 5-July-2022) analysis of the same set of contigs. These scBCR-seq annotations were merged with the Seurat data object holding scRNA-seq data as detailed above. The dataset was filtered to retain B cells fulfilling the following criteria: (1) with matching heavy/light chain sequence from scBCR-seq, and; (2) non-zero expression of both *Cd19* and *Ms4a1.* A resolution parameter of 0.4 was used in Seurat::FindClusters to determine cell clusters. sciCSR was applied to infer transitions both between cell clusters and between BCR isotypes, using (1) the signature-based CSR potential, (2) SHM level, and (3) RNA velocity information. For transitions between cell clusters, the source state was chosen to be cluster ‘6’ (naïve B cells, these cells have typically high IgM expression, see Supplementary Figure S9) and the target state as cluster ‘9’ (plasmablasts, these cells typically produce a large amount of transcripts mapping to V genes, see Supplementary Figure S9). For inferring CSR, IgM was chosen as the source and IgA as the target state.

#### SARS-CoV-2 vaccination longitudinal follow-up from Kim et al^73^

Raw FASTQ files were downloaded from SRA accession SRP356296 and aligned as described above; these alignments were used as input for sciCSR to enumerate productive/sterile transcripts. The scRNA-seq cell-by-gene count data were downloaded from the Zenodo repository (https://doi.org/10.5281/zenodo.5895181) as a python AnnData object. Only cells from donors 07 and 20 at weeks 4 (day 28), 7 (day 60), 15 (day 110) and 29 (day 201) were retained, exported as .h5ad files and converted to R Seurat object using the R SeuratDisk package (v0.0.0.9020). Since the count data were already preprocessed and normalised, this Seurat object was directly analysed. scBCR-seq data from the same cells were downloaded from the same Zenodo repository. This data table annotated heavy-chain sequences as IgM/G/A but lacked annotations of the subclasses. IgBLAST^101^ (v1.19.0) was used (with default parameters) to call subclasses for these sequences, using nucleotide sequences of germline immunoglobulin alleles downloaded from IMGT (accessed 28-July-2022) as reference. The human C-region artificially spliced exons sets were downloaded (https://www.imgt.org/vquest/refseqh.html#refdir2) and used as reference set. The scBCR-seq table annotated with heavy-chain subclass information was merged into the metadata slot of the Seurat data objects. Only the cells labelled “GC” (germinal centre) in the author-provided cell type annotation were considered. Calculation of CSR potential and SHM level was carried out using sciCSR as described above. The transition inference method in sciCSR was used using the CSR potential as pseudotime, to infer CSR using IgM as the source state and IgG4 as the target state; since there were no cells with IgE or IgA2 as their BCR isotypes, IgG4 was the state furthest along the IgH locus for this dataset. The isotype annotation was taken from the merged scBCR-seq metadata.

To compare CSR inference with observed BCR isotype distribution, we first calculated, for each isotype, the total inward flux (i.e. amount of flux towards each isotype) inferred using TPT, and reasoned that this total inward TPT flux should predict the isotype distribution observed in a subsequent timepoint. We therefore compared the TPT inward flux at week 7 to scBCR-seq isotype distribution at week 15, and TPT inward flux at week 15 to observation at week 29. The similarity of the TPT-predicted and observed distributions was evaluated using cosine similarity, calculated using the R philentropy^98^ package (v0.5.0).

### Reference atlases

We integrated previously published datasets to generate the following B cell atlas, profiled using 10X Genomics 5’ technologies, for obtaining reference isotype signatures in estimating CSR potentials for user-supplied datasets. In each case the aligned BAM files were used as input to sciCSR for quantifying productive/sterile transcripts and the resultant count matrix was decomposed using NMF to derive a reference signature matrix (see section *“Deriving CSR potential”* of Materials and Methods).

### Human B cell atlas

We integrated the following two scRNA-seq datasets of human B cells:

(1) Data from Stewart et al.^10^ containing B cells from the peripheral blood FACS-sorted into

5 phenotypically defined populations (Transitional [CD19+IgD+CD27’CD10+], Naïve [CD19+IgD+CD27^-^CD10^-^], IgM memory [CD19+IgD+CD27+], Classical Memory

[CD19+IgD’CD27+], Double Negative [CD19+IgD’CD27’]). Data from the donor HB6 were considered in this atlas. Data pre-processing was previously described^10^ and the aligned BAM files were directly taken as input to sciCSR to generate productive/sterile transcript counts, and merged with the R Seurat data object available at ArrayExpress accession E-MTAB-9544.

(2) Data from King et al.^11^ containing B cells from human tonsil samples. Raw FASTQ files were downloaded from ArrayExpress accession E-MTAB-9005 and processed as described above for sciCSR productive/sterile transcript quantification. B cell scRNA-seq transcriptomic count data were downloaded as a Seurat data object from the same ArrayExpress accession record.

### Mouse B cell atlas

We processed and integrated the following two datasets to form a mouse B cell atlas:

(1) Data from Mathew et al.^13^ for B cells from mediastinal lymph node, lung and spleen of mice at days 7, 14 and 28 days after influenza infection. Raw FASTQ files were downloaded from ArrayExpress accession E-MTAB-9478 (scRNA-seq) and E-MTAB-9491 (scBCR-seq) and pre-processed as described above using the “cellranger multi” function.

(2) Data from Luo et al.^14^ for peritoneal B cells sampled from healthy neonates, young adults, and elderly mice. Raw FASTQ files were downloaded from ArrayExpress accession E-MTAB-10081 and pre-processed using “cellranger multi”. Only data from samples D, E and F were considered as scBCR-seq and scRNA-seq data were obtained in parallel only for these samples.

### Data integration

For the mouse atlas count matrices from Mathew et al. and Luo et al. were read in and directly combined prior to normalization, dimensionality reduction and clustering. For the human atlas, the two Seurat data objects holding data from King et al. and Stewart et al. were integrated by following the data integration protocol in the Seurat^71^ package. Briefly, anchoring points for data integration were established using the SelectIntegrationFeatures followed by FindIntegrationAnchors function, and the two datasets were integrated using the IntegrateData function in Seurat. All functions were evaluated using default parameters.

