## Supplementary Materials for "sciCSR infers B cell state transition and predicts class-switch recombination dynamics using single-cell transcriptomic data"

### Supplementary Note

#### Supplementary Note 1. Robustness of sciCSR in studying rare populations.

Here we investigate the minimum number of cells needed for sciCSR to accurately recover transitions underlying the data. Recall from the manuscript & Methods that sciCSR provides CSR/SHM potential scores for CellRank[1] to fit Markov State Models (MSM). Intuitively, the number of available observations would affect the ability of MSM to correctly identify states and infer transitions between them. One way to answer this question by generating a synthetic dataset of size comparable to test cases presented where sciCSR successfully recovers transitions, and apply sciCSR on this synthetic dataset. By downsampling the number of cells in each state and re-apply sciCSR on this data, we can then compare the inference results to the ground-truth.

*Description of synthetic dataset*

We consider a simple mixture of three states: IgM, IgG1 and IgA1, and the class-switching dynamics between these states. This can be taken as the transitions between three B cell subsets: Naïve, Classical Memory, DN1, as we found previously that these B cell subsets are enriched respectively in IgM/IgG1/IgA1 expression[2] (Stewart et al). For simplicity we set each state to have an equal number ($n$) of cells, and calculate CSR potential as input for sciCSR. $n$ is the only variable in the simulation.

We require two inputs: (1) transcriptomic gene counts to build k-nearest neighbour (kNN) graph to describe the structure of the data, and (2) CSR/SHM potentials as pseudotime ordering to bias the kNN graph and describe the transitions. Here we used the splatter[3] package to simulate a count matrix of 2,000 genes for a three-group (or ‘state’) mixture with 10,000 cells in each state to constitute the ground-truth; the Stewart et al[2] dataset of healthy volunteer B cells in circulation was used as input such that the splatter-simulated gene counts would fit to a distribution similar to the Stewart et al. data. The splatter-simulated gene counts served to build the kNN graph. To assign the CSR potential, we randomly sampled the NMF-decomposed weights from Stewart et al and assign to each state, as follows:

- Cells in the IgM state were randomly assigned CSR potential of a Naïve cell from Stewart et al.;
- Cells in the IgG1 state were randomly assigned CSR potential of a C-mem cell, and;
- Cells in the IgA1 state were randomly assigned CSR potential of a DN1 cell.

As mentioned above, these cell subsets were characterised by enrichment of these isotypes in scRNA-seq data[2]. sciCSR was invoked with the CSR potential and the kNN graph as inputs with default parameters. The resulting fluxes were compared against the ground-truth. In this simulation, with an equal number of cells in each state we would expect that the majority (~ 50%) of the total flux to be attributed to transitions from IgM to IgA1 directly, and the remaining flux separated equally between IgM → IgG1 and IgG1 → IgA1. Figure i below demonstrates that sciCSR inference recapitulates this expected outcome, and the inference is robust across a range of cell population sizes, up to groups of $n$ = 10 cells.

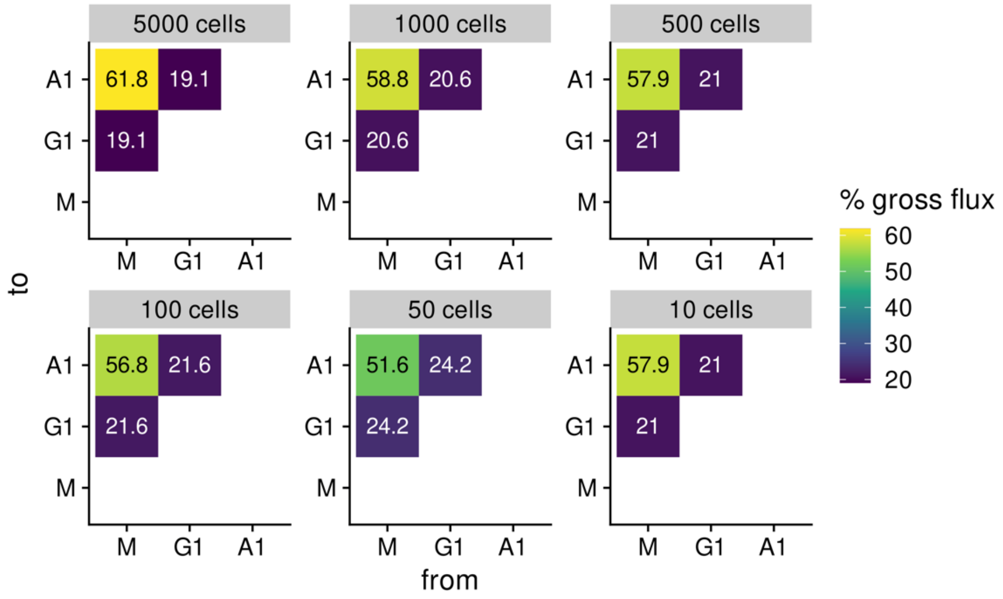


Figure i. sciCSR inference on a simulated mixture of IgM, IgG1 and IgA1 cells, with equal numbers of cells belonging to each isotype. Data were downsampled to the stated number of cells per group before inference using sciCSR. Heat colour corresponds to the proportion of flux attributed to each transition.

### Supplementary Figures

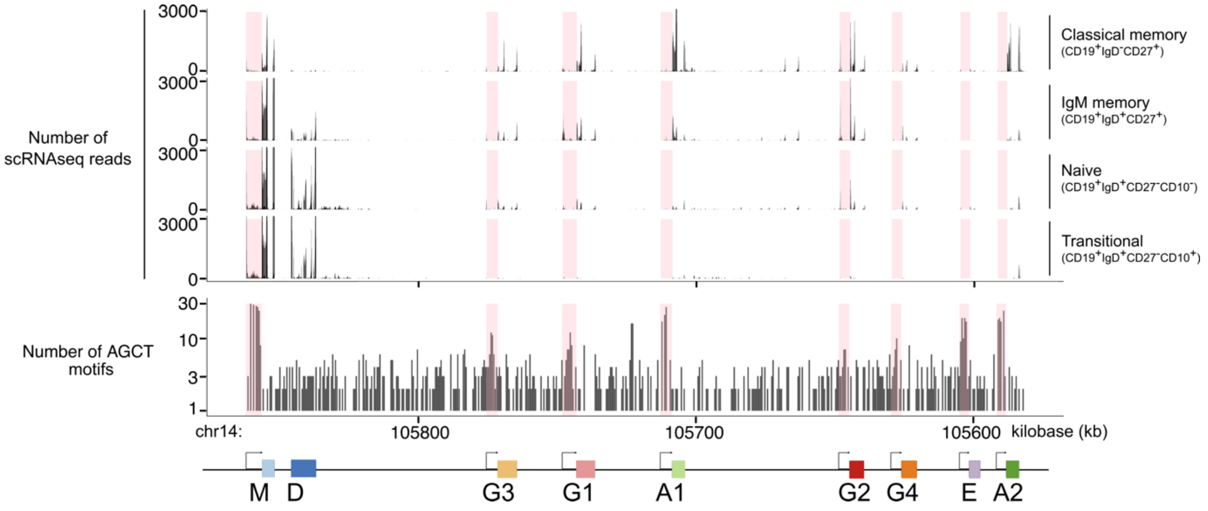


Supplementary Figure S1. Histogram of scRNA-seq read counts from the Stewart et al. dataset across the *IGH* genomic locus. The distribution of constant region genes was illustrated at the bottom of the histogram (coloured boxes) along with counts of 5'-AGCT-3' motifs in sliding windows of 500 base-pairs (histogram in the middle). Regions 5' of the constant region coding segment were highlighted with pink rectangles.

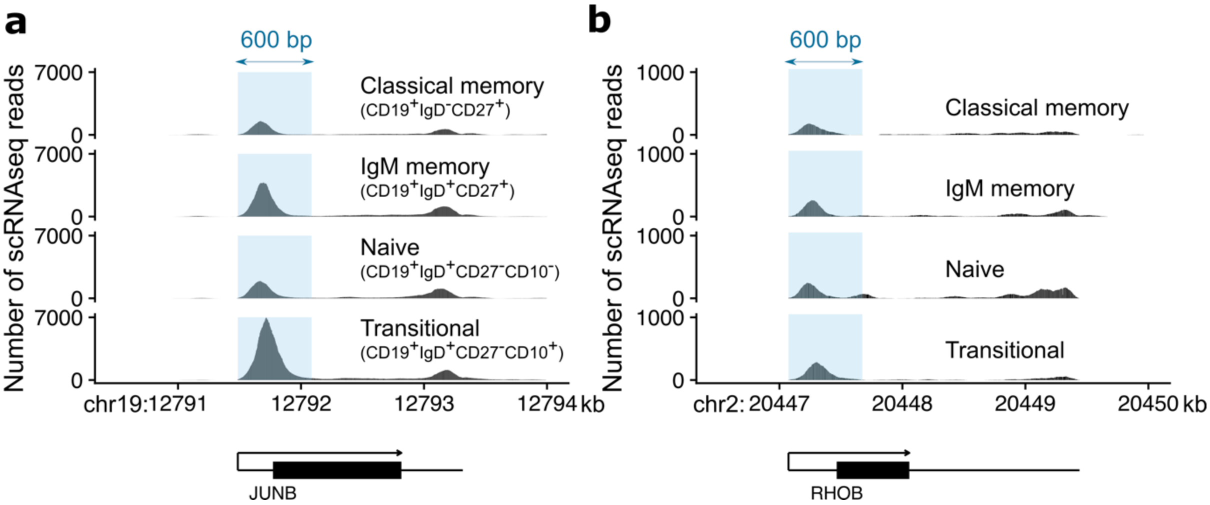


Supplementary Figure S2. Distribution of scRNA-seq reads from Stewart et al. across two single-exon transcripts *JUNB* (Ensembl transcript ENST00000302754.6, panel a) and *RHOB* (ENST00000272233.6, panel b). Regions of 600 base-pairs (bp) at the 5' end of the transcript were highlighted (blue rectangles) to illustrate the concentration of reads at this region from the scRNA-seq experiments.

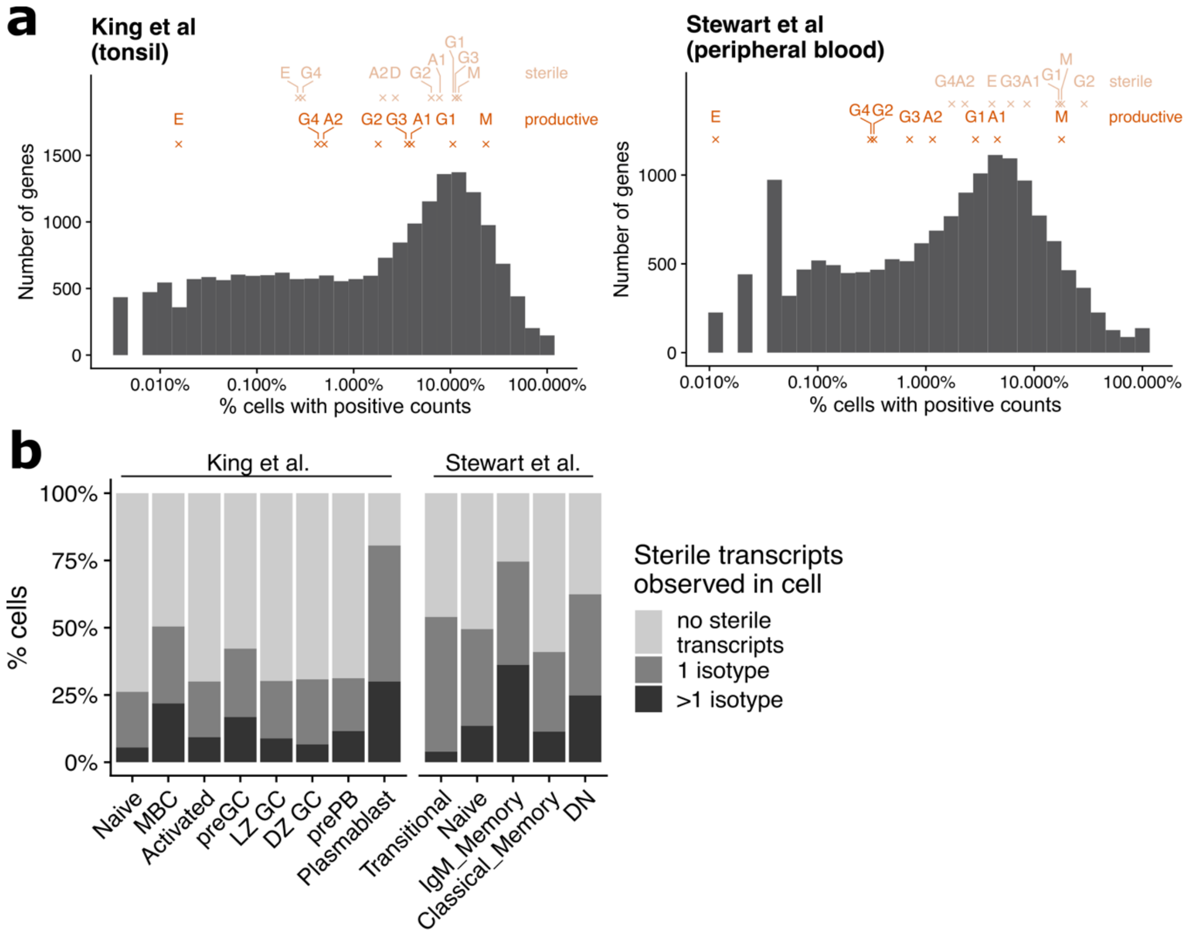


Supplementary Figure S3. Distribution of productive/sterile IgH transcripts.

(a) Sparsity of productive/sterile IgH transcripts. Grey bars depict distribution of all transcripts with count data in the King et al. tonsil B cell dataset (left) and the Stewart et al. peripheral B cell dataset (right), showing for each transcript the proportion of cells in the data with positive counts. The values for each productive and sterile IgH transcript are noted at the top of the histogram with crosses.

(b) The number of isotypes presented in the sterile transcripts observed in each cell in the King et al. and Stewart et al. datasets, expressed per cell type. Each cell was assessed in terms of the number of isotypes represented in the sterile transcripts associated with each cell.

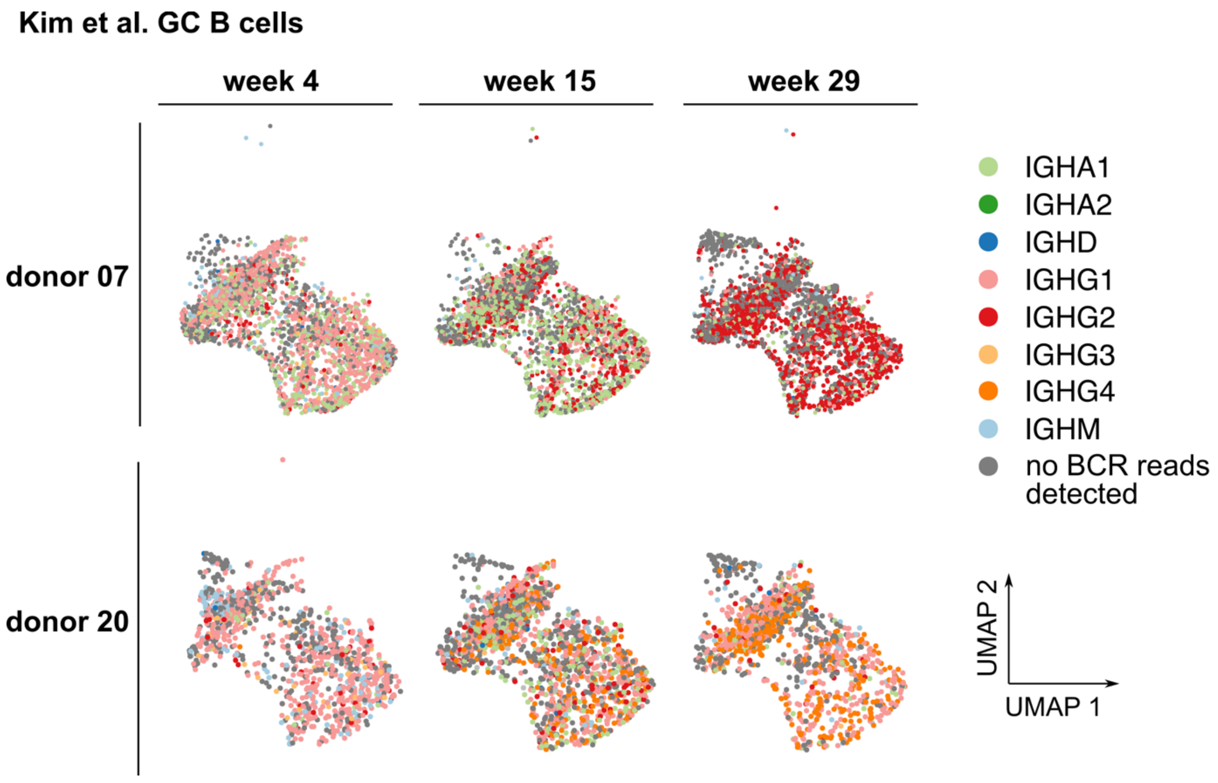


Supplementary Figure S4. UMAP visualisation of GC B cell scRNA-seq data of two donors from Kim et al., separated by timepoints (columns) and coloured by BCR isotype indicated in the single-cell-matched scBCR-seq data. All donor/timepoint combinations were projected onto the same UMAP reduction.

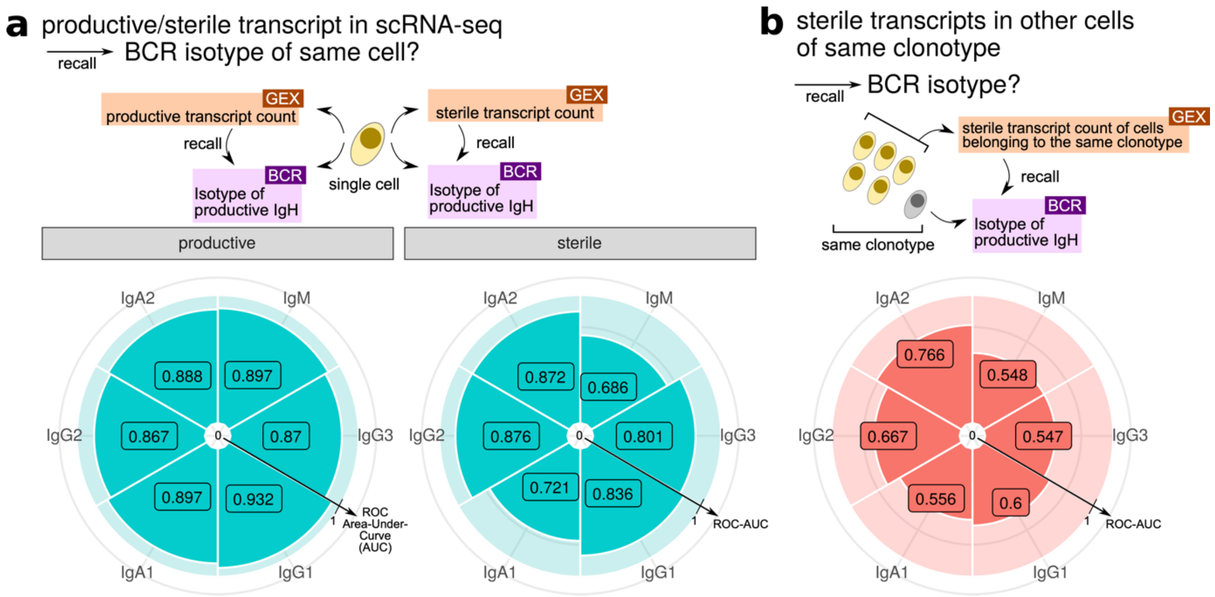


Supplementary Figure S5. Recall BCR isotypes using productive/sterile transcript count recovered using sciCSR in the King et al. tonsil B cell dataset.

(a) *(left) Productive* transcript count was recovered from scRNA-seq data using sciCSR and treated as a predictor of BCR isotype of the same cell according to the scBCR-seq data. Pizza charts depict the area-under-curve of the receiver operating characteristic (AUC-ROC) curve separately for each isotype.
*(right) Sterile* transcript count was recovered from scRNA-seq data using sciCSR and treated as a predictor of BCR isotype of the *same cell*. Pizza charts depict AUC-ROC separately for each isotype.

(b) *Sterile* transcript count was recovered from scRNA-seq data using sciCSR and treated as a predictor of BCR isotype of *cells belonging to the same clonotype*. Pizza charts depict AUC-ROC separately for each isotype.

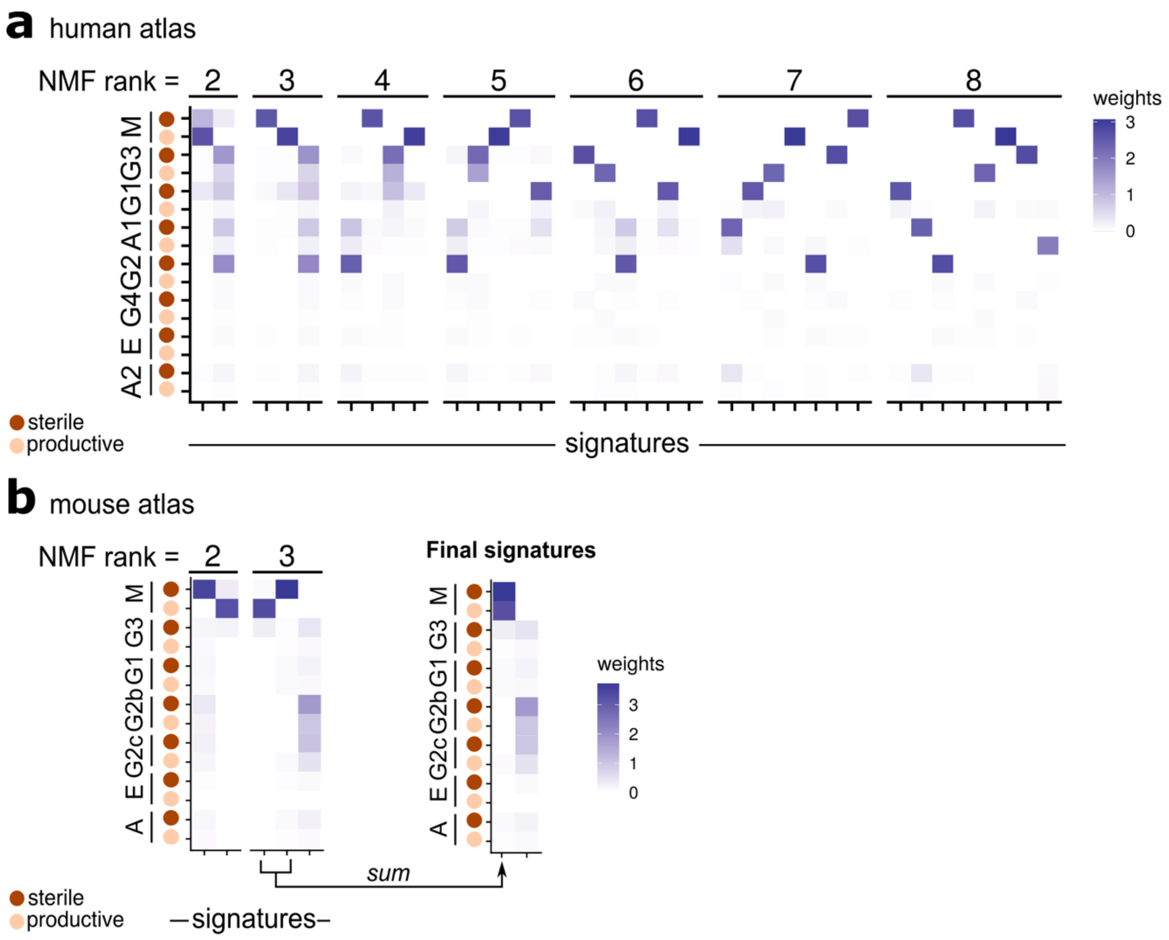


Supplementary Figure S6. Isotype signatures identified using nonnegative matrix factorization (NMF) on human and mouse B cell atlases.

(a) Effect of changing NMF rank (i.e. the desired number of signatures) on the human B cell atlas. The signature matrix under each NMF setup was shown. The final isotype signature matrix for human is the rank = 2 condition.

(b) Deriving the mouse signature matrix. (left) the NMF signature matrices using rank = 2 and 3. Neither setup derived a signature matrix similar to the rank = 2 condition in the human atlas (see panel a), although it appears that the IgM-dominant signature was separated into two signatures in the rank = 3 condition (first two columns). (right) The final signature for mouse was derived by summing together the first two signatures in the rank = 3 condition. The final signature matrix was shown here.

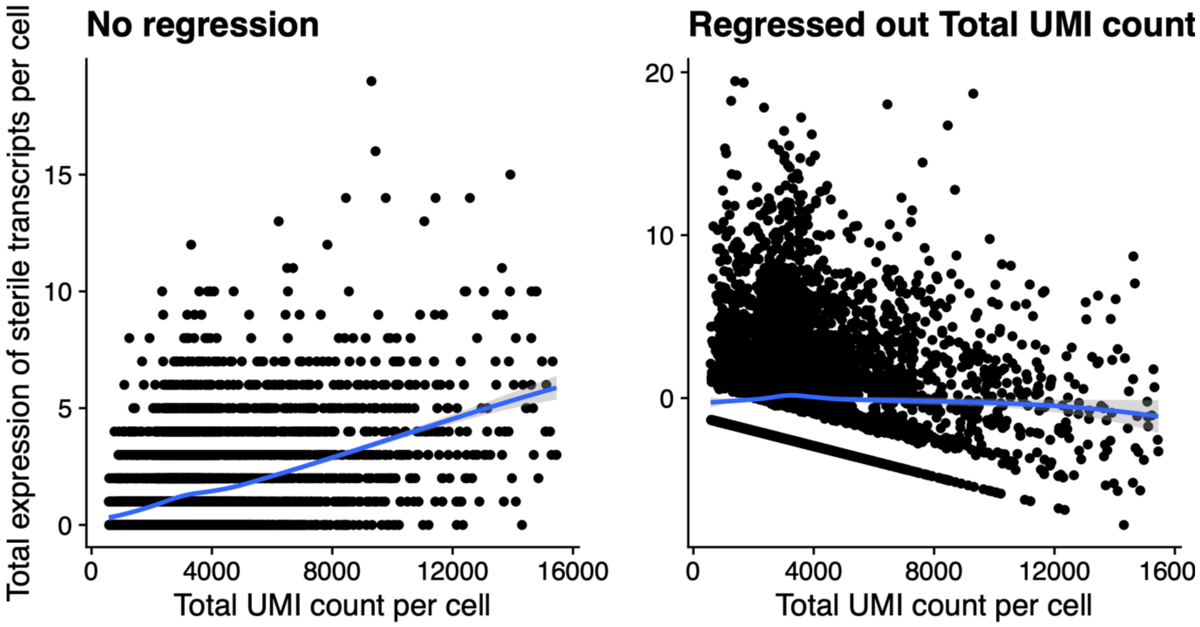


Supplementary Figure S7. Effect of total UMI count on sterile transcript expression level. The sterile transcript expression level per cell (vertical axis) was plotted as a function of the total UMI count per cell (horizontal axis), before (left panel) and after (right) regression to remove the effect of total UMI count. Data shown corresponds to the Stewart et al. dataset. Trendline was fitted using local polynomial regression (loess) fitting in R.

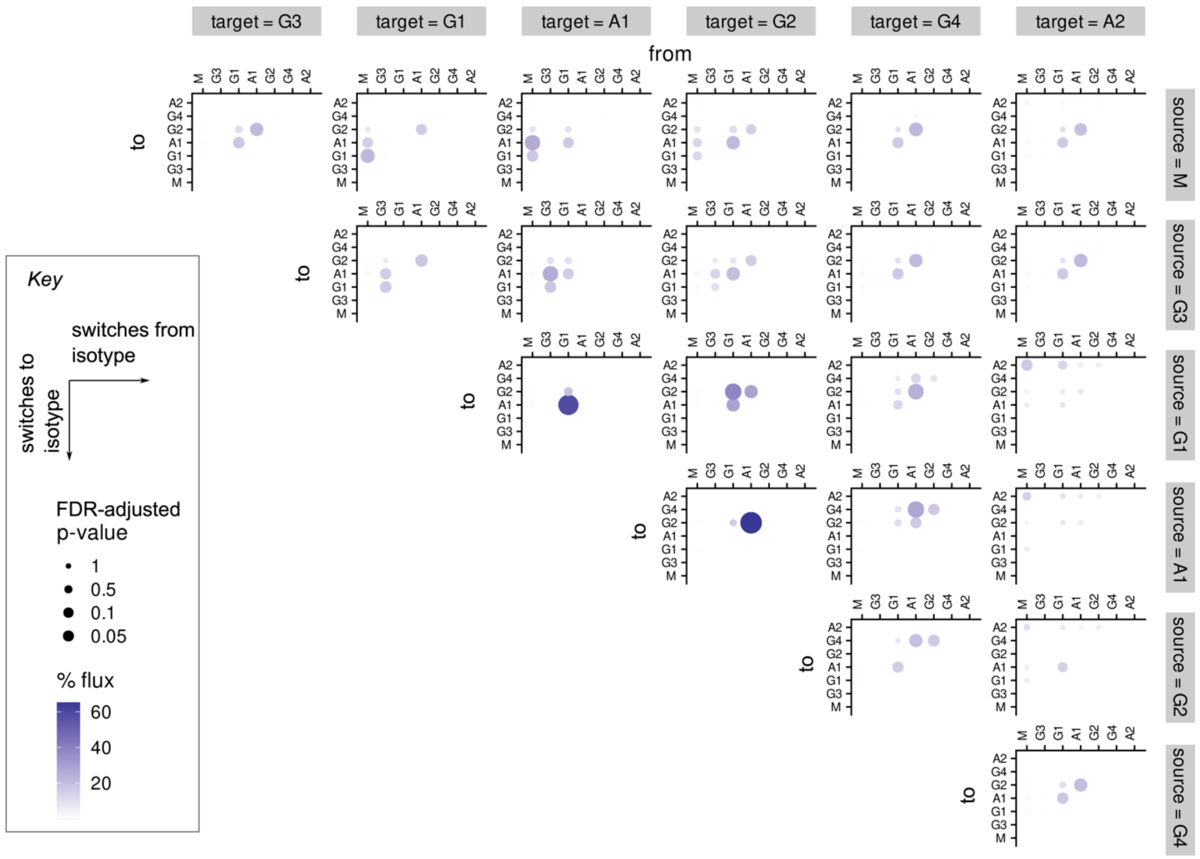


Supplementary Figure S8. Effect of selecting different source (row) and target (column) states on TPT results using the week 15 timepoint of donor 07 from the Kim et al. vaccination time-course dataset. All inferences were performed using the NMF-derived CSR potential as pseudotime input to sciCSR which was invoked using identical and default parameters. The inferred fluxes tend to be higher for transitions involving states which are either the chosen source or target states.

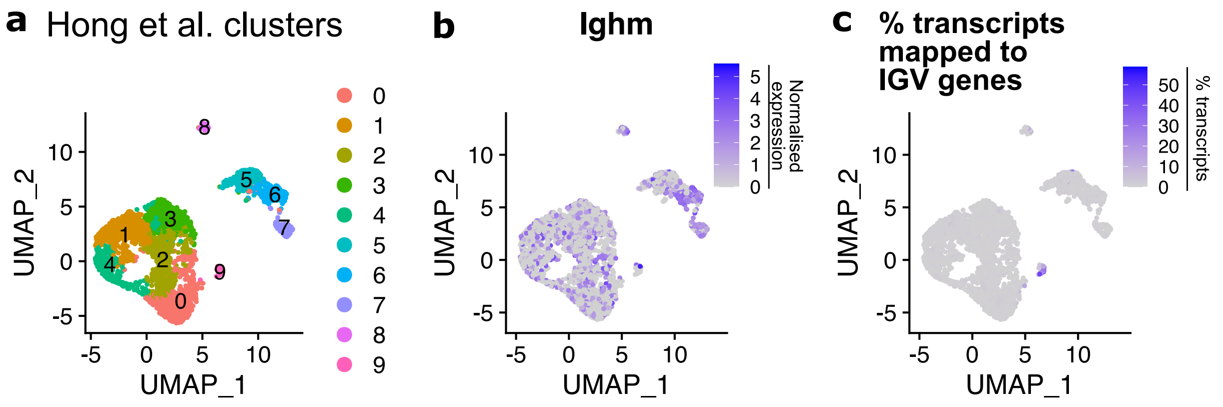


Supplementary Figure S9. Cell clusters from the Hong et al. dataset.

(a) UMAP projection of the Hong et al. data grouped into 10 cell clusters (colours).

(b) Expression of *Ighm* gene in the Hong et al. data.

(c) Percentage of transcripts which are mapped to IGV genes for each cell expressed as a heat scale.

### Supplementary Tables

**Supplementary Table 1. Genomic coordinates of human and mouse constant region genes and their corresponding sterile transcript start sites.** Sterile transcript start sites were mapped by aligning collected sterile transcript sequences from the literature to the hg38 (human) and mm10 (mouse) reference genome assemblies (see Methods). Note that the IgH genes are located on the minus strand for both mouse and human. This table is shipped with the sciCSR R package.

| Ensembl gene ID | Gene | Genomic coordinate | Sterile transcript start site | Genome assembly | Organism |
|---|---|---|---|---|---|
| ENSG00000211899 | *IGHM* | chr14:105,851,705-105,856,218 | 105,861,958 | hg38 | Human |
| ENSG00000211897 | *IGHG3* | chr14:105,764,503-105,771,405 | 105,775,495 | hg38 | Human |
| ENSG00000211896 | *IGHG1* | chr14:105,736,343-105,743,071 | 105,748,177 | hg38 | Human |
| ENSG00000211895 | *IGHA1* | chr14:105,703,995-105,708,665 | 105,712,867 | hg38 | Human |
| ENSG00000211893 | *IGHG2* | chr14:105,639,559-105,644,790 | 105,648,651 | hg38 | Human |
| ENSG00000211892 | *IGHG4* | chr14:105,620,506-105,626,066 | 105,629,851 | hg38 | Human |
| ENSG00000211891 | *IGHE* | chr14:105,597,691-105,601,728 | 105,605,176 | hg38 | Human |
| ENSG00000211890 | *IGHA2* | chr14:105,583,731-105,588,395 | 105,591,896 | hg38 | Human |
| ENSMUSG00000076617 | *Ighm* | chr12:113,418,558-113,422,730 | 113,427,389 | mm10 | Mouse |
| ENSMUSG00000076615 | *Ighg3* | chr12:113,356,224-113,361,232 | 113,366,394 | mm10 | Mouse |
| ENSMUSG00000076614 | *Ighg1* | chr12:113,325,240-113,330,523 | 113,339,379 | mm10 | Mouse |
| ENSMUSG00000076613 | *Ighg2b* | chr12:113,302,965-113,307,933 | 113,314,819 | mm10 | Mouse |
| ENSMUSG00000076612 | *Ighg2c* | chr12:113,285,325-113,288,932 | 113,296,262 | mm10 | Mouse |
| ENSMUSG00000087642 | *Ighe* | chr12:113,269,260-113,273,248 | 113,277,549 | mm10 | Mouse |
| ENSMUSG00000095079 | *Igha* | chr12:113,254,830-113,260,236 | 113,265,138 | mm10 | Mouse |

### Supplementary Data

**Supplementary Data 1. Nucleotide sequences input for generating simulated RNA-seq reads using the polyester software.** (FASTA file)
